# Tripartite ER-Mitochondria-Lipid Droplets contact sites control adipocyte metabolic flexibility

**DOI:** 10.1101/2025.09.16.676493

**Authors:** Marie Palard, Gilliane Chadeuf, Maxime Carpentier, Samuel Frey, Mikael Croyal, Rola Shaaban, Samy Hadjadj, Chloe Cloteau, Naima El Khallouki, Perrine Bomme, Soazig Le Lay, Manuel Marjouh, Perrine Paul-Gilloteaux, Alexandre Humbert, Christine Durand, Nicolas Bertocchini, Yoann Combot, Antria Antreou, Cedric Le May, Simon Ducheix, Bertrand Cariou, Francesca Giordano, Jennifer Rieusset, Abdou Rachid Thiam, Xavier Prieur

**Affiliations:** Université de Nantes, CNRS, INSERM, l’institut du thorax, 44000 Nantes, France; Laboratoire de Physique de l’École normale supérieure, ENS, Université PSL, CNRS, Sorbonne Université, Université Paris Cité, F-75005 Paris, France; Nantes Université, CNRS, CHU Nantes, INSERM, l’institut du thorax, F-44000, Nantes, France; Nantes Université, CHU Nantes, INSERM, CNRS, SFR Santé, Inserm UMS 016, Nantes, France; Université Paris-Saclay, CEA, CNRS, Institute for Integrative Biology of the Cell (I2BC), 91190 Gif-sur-Yvette, France; Ultrastructural BioImaging Core Facility, Department of Cell Biology and Infection, C2RT, Institut Pasteur, Université Paris Cité, Paris, France; Univ Angers, SFR ICAT, 49000 Angers, France; Nantes Université, CHU Nantes, CNRS, Inserm, BioCore, US16, SFR Bonamy, Nantes, France; Laboratoire CarMeN, UMR INSERM U1060/INRA U1397, Université Claude Bernard Lyon1, Pierre-Bénite, France

**Author notes:** shared last authors. **Corresponding authors** Jennifer Rieusset, Abdou Rachid Thiam, Xavier Prieur.

## Abstract

Obesity is a major risk factor for cardiometabolic diseases, with adipocyte dysfunction playing a central role. Understanding how lipid storage and mobilization are regulated—and disrupted—in adipocytes is key to addressing obesity-associated complications. The ER-anchored protein Seipin controls lipid droplet (LD) biogenesis and maintenance, and its loss disrupts ER–LD contact sites. In humans, Seipin deficiency causes generalized lipodystrophy, a severe form of adipocyte dysfunction. We previously showed that Seipin also localizes at ER–mitochondria contact sites (MAM), where it regulates calcium exchange and mitochondrial function. Here, we examined whether Seipin targeting to MAM and ER–LD sites overlaps functionally. We analyzed subcutaneous adipose tissue (AT) from inducible Seipin-knockout mice using transmission electron microscopy (TEM) and proximity ligation assays (PLA) to quantify membrane contact sites (MCS) involving the ER, LDs, and mitochondria. In control mice, feeding reduced MAMs while increasing ER–LD and mitochondria–LD contacts, whereas Seipin deficiency abolished this remodeling. Specifically, under lipid loading, MAMs located in proximity to LDs—tripartite contact sites known as MAM–LD—were increased in control but not in Seipin-deficient adipocytes. Fluorescence recovery after photobleaching assays revealed that Seipin depletion impairs triglyceride transfer to LDs, an effect rescued by the MAM–LD–reinforcing synthetic peptide Linker-ER-Mi. Importantly, this rescue was abolished by silencing the mitochondrial calcium uniporter, demonstrating that calcium exchange is critical for triglyceride storage in LDs. We further investigated how MAM–LD remodeling influences adipocyte metabolic flexibility. Using TEM and PLA, we monitored two MAM subtypes: those forming MAM–LD and those engaging cytosolic mitochondria (MAM–CM). During adipogenesis, MAM–LD frequency increased while MAM–CM decreased. Similarly, in mouse AT and 3T3-L1 adipocytes, lipid loading selectively promoted MAM–LD. Notably, this adaptive remodeling of membrane contact sites was blunted in the adipose tissue of diet-induced obese mice. Genetic disruption of MCS in 3T3-L1 adipocytes altered lipid flux, impaired lipolysis, and reduced insulin signaling. Together, our findings identify MAM–LD contacts as key regulators of adipocyte lipid handling and metabolic flexibility, whose disruption may underlie the metabolic inflexibility of obesity.

## Introduction

The increasing prevalence of obesity is driving a rise in cardiometabolic morbidities, with adipocyte dysfunction playing a central role^1^. In obese individuals, adipose tissue (AT) exhibits significant impairments in both lipid storage and mobilization. Stable isotope tracer studies show that, under normal conditions, lean individuals efficiently store lipids postprandially and release them during fasting. In contrast, these processes are markedly impaired in individuals with obesity, reflecting a profound metabolic inflexibility^2^. Similarly, the use of atmospheric ^14^C incorporation into lipids reveals decreased lipid turnover in obese subjects^3^. These findings suggest that in obesity, AT enters a state of metabolic inertia, reducing its capacity to store excess lipids and promoting ectopic lipid accumulation in non-adipose tissues—thereby contributing to cardiometabolic complications^4^. Furthermore, recent studies indicate that even after weight loss, AT retains features of dysfunction that predispose to weight regain and exacerbate metabolic complications^5^. This highlights the urgent need to better understand the mechanisms governing lipid storage and mobilization in adipocytes and how these processes are dysregulated in obesity.

Congenital generalized lipodystrophy (CGL) represents the most severe form of primary adipocyte dysfunction, characterized by a near-complete absence of AT and severe metabolic complications^6^. Bi-allelic mutations in the BSCL2 gene, which encodes the endoplasmic reticulum (ER)-anchored protein seipin, are the most common cause of CGL^7^. Seipin deficient mice display severe lipoatrophy and extreme insulin resistance^8^. Furthermore, inducible seipin deficiency leads to rapid AT loss^9^. Given the severity of the phenotype associated with seipin deficiency, we and others hypothesize that elucidating seipin’s function could reveal novel pathways that regulate adipocyte homeostasis^8^. Early studies in yeast demonstrated that seipin deficiency leads to abnormal lipid droplet (LD) morphology^10,11^. Seipin is essential for both the initiation of LD biogenesis^12^ and the transition to mature LDs ^13^. Seipin is enriched at ER–LD contact sites, and in fibroblasts derived from patients with seipin deficiency, these contacts appear disorganized ^14^. In addition, we have shown previously that, in adipocytes, seipin is enriched at ER–mitochondria contact sites (also known as mitochondria-associated membranes, or MAMs), and that seipin loss disrupted ER-to-mitochondria calcium flux^15^. Importantly, in fed conditions, seipin is enriched in ER-LD contacts, whereas starvation promotes its relocalization to MAMs. Whether seipin recruitment to MAMs and ER–LD contact sites is mutually exclusive remains unknown. Additionally, the function of seipin at MAMs is not well understood, and it is unclear whether seipin localized at MAMs contributes to its central role in regulating lipid storage in adipocytes.

In the past decade, membrane contact sites (MCS) have emerged as central regulators of cellular metabolic flexibility, i.e., the capacity to adapt fuel utilization to nutritional and hormonal states^16^. Among these, MAMs serve as key hubs for lipid and calcium exchange. By integrating nutrient and hormonal cues, MAMs modulate both insulin sensitivity and metabolic flexibility ^17,18^. In the liver, disruption of MAMs promotes insulin resistance and metabolic dysfunction such as MASLD, whereas reinforcement of these contacts protects high-fat diet (HFD)-fed mice from metabolic complications^19^. ER–LD contact sites are also critical for lipid storage, and their perturbation has been linked to various metabolic diseases ^20^. Indeed, LDs originate from the ER^21^ and remains physically contiguous with it to exchange proteins and lipids and can also establish non-contiguous contacts with the ER throughout their lifetime. More recently, mitochondria–LD (Mi–LD) contact sites emerged as essential in lipid homeostasis for both anabolic and catabolic processes. In brown AT, two distinct mitochondrial subpopulations have been described: cytosolic mitochondria (CM) and peri-droplet mitochondria (PDM) ^22^, which we call here MAM-LD. In the liver, PDM are thought to promote lipogenesis, while CM primarily support catabolic processes^23^. This subcellular heterogeneity also complexifies our understanding of MAMs: MAMs involving CM would be the “classical” MAMs, known to support catabolic processes during fasting^17^, while those involving PDM (referred to as MAM–LDs) are enriched in lipid synthesis enzymes^23^. To date, very few reports have focused on MAM-LD, and to our knowledge, none have examined these in adipocytes.

In this study, we seek to verify the hypothesis that alteration of organelle contact sites could contribute to adipocyte dysfunction and thus be a new molecular mechanism underlying cardiometabolic complications associated with obesity and lipodystrophies. We firstly characterized the MCS in seipin deficient adipocytes, and we assessed their functional involvement in lipid handling. Then, we characterize adipocyte’s MCS of the most common form of adipocyte dysfunction by analysing the AT of obese mice. Finally, *in vitro*, we used several genetic interventions to specifically assess the consequences of MCS disturbances on adipocyte homeostasis.

## Results

### Seipin deficiency alters MAM-LD contacts and abolishes the nutrient-driven organelles remodeling *in vivo* in mice

We previously demonstrated that seipin localizes in the MAM^15,24^, which led us to investigate here whether seipin deficiency affects MAMs. We analyzed subcutaneous AT from inducible adipocyte-specific seipin KO mice (iATSKO) 15 days post-tamoxifen injection. At this stage, AT displays only mild dysfunction, enabling us to capture the early mechanisms leading to 80% adipose mass loss observed in 3-month-old iATSKO mice. We used transmission electron microscopy (TEM) to quantify changes in inter-organelle contacts (Figure 1A). Specifically, we assessed the total number of mitochondria-ER contacts (i.e., Total MAMs), normalized to mitochondrial area. Further, we classified them as either associated with LDs (i.e., MAM–LD) or independent of LDs, which has been previously called MAM-CM^23^, involving cytosolic mitochondria (CM). We also quantified ER–LD and mitochondria–LD (Mi–LD) contacts and measured the extent of contact between organelles. Our analysis revealed a ∼25% reduction in total MAMs (Figure 1B). Functionally, seipin-deficient AT mitochondria exhibited reduced oxygen consumption and structural abnormalities (Supplementary Figure 1C). In control mice, mitochondria engaged with lipid droplets (peridroplet mitochondria, PDM) were consistently longer and broader than cytosolic mitochondria (CM), a distinction that was lost in iATSKO adipocytes (Supplementary Figures 1A–B). A global assessment of contact site lengths further revealed marked reductions in MAM, ER–LD, and Mi–LD contacts, with Mi–LD contacts showing the most substantial decrease (∼70%) (Figure 1C). The surface of these contacts was also reduced in seipin absence (Supplementary 1 D-F).

**Figure 1.**
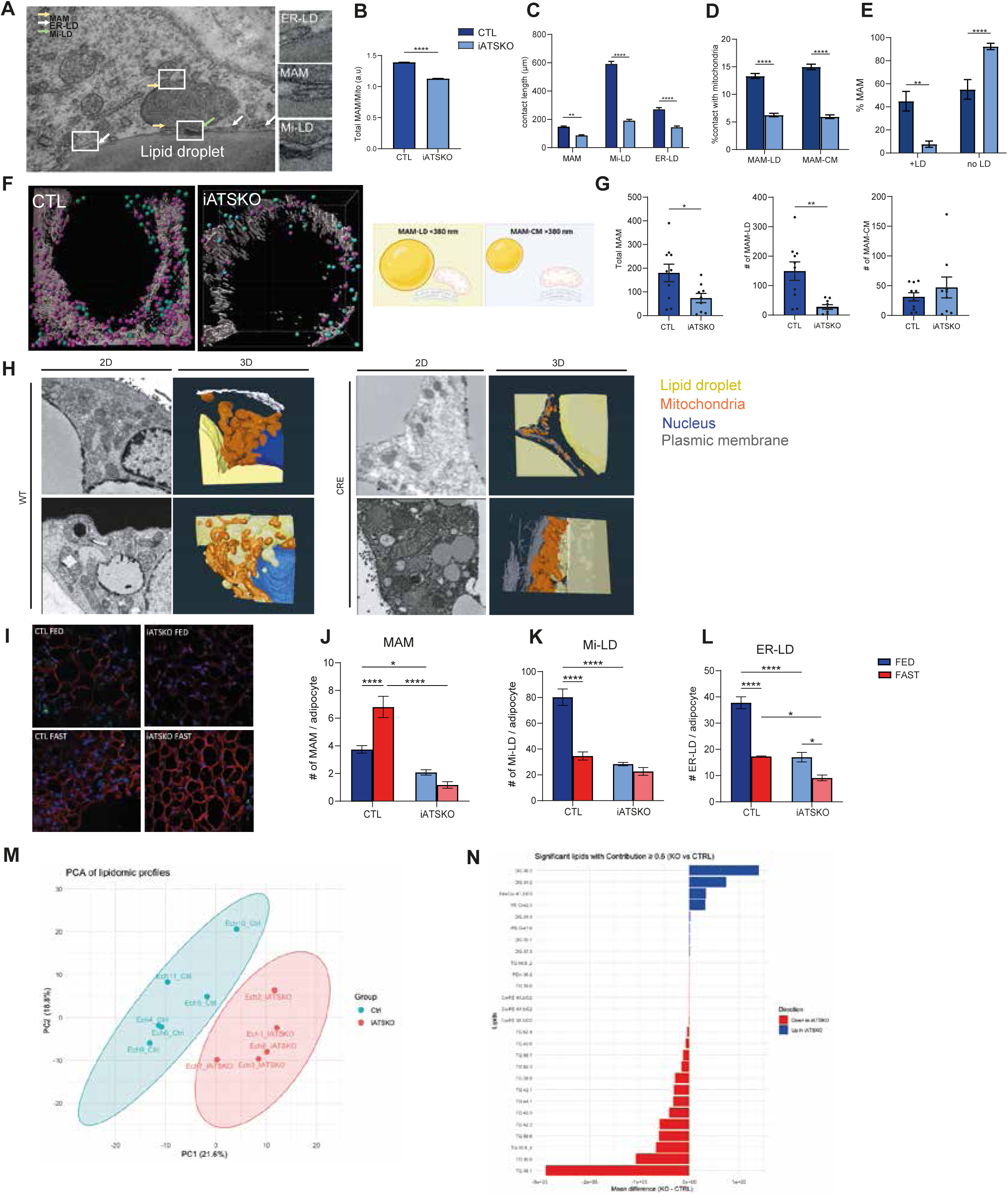
MAM-LD architecture is strongly altered in inguinal AT of Seipin-deficient mice. Inguinal adipose tissue was collected from *Bscl2^lox/lox^*(Control) and *Bscl2 ^lox/lox^ERT2-Adipoq-Cre* (iATSKO) mice in the random-fed state, 14 days after tamoxifen injection (100 mg/kg). The number of mitochondria-associated membranes (MAMs) (A), were analyzed by transmission electron microscopy (TEM). The lengths of ER–mitochondria (MAM), mitochondria–LD (Mi–LD), and ER–LD contact sites were quantified using FIJI (B). MAM-LD and MAM-CM were quantified using Fiji (C-E). Bars: mean ± SEM, *n* = 2 mice per group, 5 cells per mouse. *p* < 0.05, **p** < 0.001, ***p*** < 0.0001, Mann–Whitney test. The number of mitochondria-associated membranes (MAMs) was quantified using a proximity ligation assay (PLA) for IP3R and VDAC, combined with PLIN1 immunofluorescence, and analyzed by Structured Illumination Microscopy (F-G). Representative images show PLIN1 in grey, MAM-LD in magenta, and MAM-CM in pink (F). Total PLA dots were analyzed, and further distinguished by their distance from lipid droplets (LD): MAM-LD (<380 nm from LD) and MAM-CM (>380 nm from LD). Representative images of FIB-SEM acquisitions on adipose tissue sections from CTL and iATSKO mice (H). Cryopreserved subcutaneous adipose tissue from control and iATSKO mice in the fed or 18-hour fasted state was analyzed by immunofluorescence for PLIN1 (I) to assess tissue quality. Proximity ligation assays (PLA) quantified the following contact sites: ER–mitochondria (VDAC/IP3R; J), mitochondria–LD (PTPIP51/PLIN1; K), and ER–LD (VAPB/PLIN1; L) Bars: mean ± SEM, *n* = 5 mice per group, >100 cells per mouse. *p* < 0.05, **p** < 0.001, ***p*** < 0.0001, Mann–Whitney test. LD were isolated from iATSKO and control mice, and lipidomic profiles were performed. PCA analysis are represented (M) and bar plots highlight the lipids that contribute the most to the separation (N).

Given the reported functional differences observed between MAM-LD and MAM-CM^23^, we quantified these subtypes. Seipin deficiency reduced the surface areas of both MAM-CM and MAM-LD (Figure 1D). However, while control mice showed comparable MAM-CM and MAM-LD levels, iATSKO mice exhibited ∼90% MAM-CM dominance, with MAM-LD contacts decreasing to ∼10% (Figure 1E).

To further assess the analysis of MAM-LD at the three-dimensional level, we combined *in situ* proximity ligation assays (PLA) with immunofluorescence and structured illumination microscopy (SIM) to segment signals by LD proximity. Following a validated protocol^25^, we classified ER-Mito contacts as MAM-LD when their distance to LD was below 380 nm; otherwise, we considered them as basic MAM. Seipin-deficient AT showed a marked reduction in MAM-LD, while MAM was not significantly different between genotypes (Figure 1F-G). We used FIB-SEM (Focused Ion Beam-Scanning Electron Microscopy) to have a better visualization of the three-dimensional organization of the organelles. In samples from control mice, lipid droplets and mitochondria are closely apposed, indicating a tight spatial association. By contrast, in seipin-deficient mice, mitochondria appear detached from lipid droplets and display a more disorganized arrangement. In these datasets, ER segmentation was not feasible due to insufficient resolution. Qualitatively, these observations support the notion that seipin loss leads to a disruption of organelle spatial organization. (Figure 1H, animation Supplemental video 1 and 2).

Next, we probed the impact of the nutritional state on the remodeling of these contacts. Nutritional-state comparisons using PLA revealed that feeding reduced Total-MAMs (IP3R–VDAC) but increased ER–LD (VAPB–PLIN1) and Mi–LD (PTPIP51–PLIN1) contacts in control mice. This remodeling of contact sites was abolished in iATSKO mice, except for ER-LD, although the extent of the decrease was lower than in control mice (Figure 1I–L).

To assess LD property changes, we performed a gentle isolation of LDs (3000 g) to preserve PDM and MAM-LD together with LD, in the lipid floating fraction. Proteomic analysis detected mitochondrial markers but no genotype-dependent proteome changes (Supplemental 1H). Lipidomic analysis, however, showed strong iATSKO/control separation (Figure 1M) driven by decreased triglycerides (TG) and increased diglycerides (DG), suggesting defective TG synthesis/targeting (Figure 1N). Predictive analysis confirmed TG decrease and DG increase as genotype predictors (Supplemental 1I). Notably, at this stage, LD surface area assessed by TEM was not altered in the absence of seipin (Supplemental 1G).

Collectively these results show that seipin deficiency impairs nutritional remodeling of MCS, particularly MAM-LD structure, and associates with reduced LD TG storage and DG accumulation.

### In seipin deficient adipocytes, MAM-LD alteration alters TG flux toward LDs

Seipin prevents TG from escaping LDs ^26^ and localizes at MAMs. Thus, the reduction of organelle contacts in seipin-deficient conditions, particularly MAM–LD contacts in adipocytes, led us to investigate whether seipin, or seipin together with MAM-LD, is required for efficient lipid storage into LDs. To address this, we used 3T3-L1 adipocytes transfected with either non-targeting control siRNA (siNTC) or BSCL2 siRNA (siBSCL2). TEM analysis confirmed that siBSCL2 treatment reproduced the MCS defects observed in iATSKO mice, showing decreased ER-mitochondrial, ER–LD, and Mi-LD contacts under oleic acid (OA) loading (Figure 2A-B). In control cells, lipid loading reduced overall MAM surface area but concomitantly increased MAM-LD surface area, a remodeling response that was absent in siBSCL2-treated adipocytes (Figure 2C-E). FIB-SEM imaging combined with organelle segmentation further revealed that, in control cells, the three organelles are tightly arranged, with ER–mitochondria–LD and mitochondria–ER–LD configurations being frequently observed, supporting the existence of stable tripartite contacts. In contrast, in seipin-deficient cells, both mitochondria and ER appear loosely organized around lipid droplets, indicating a loss of this tight spatial integration. (Figure 2-F and supplemental video 3 and 4).

**Figure 2:**
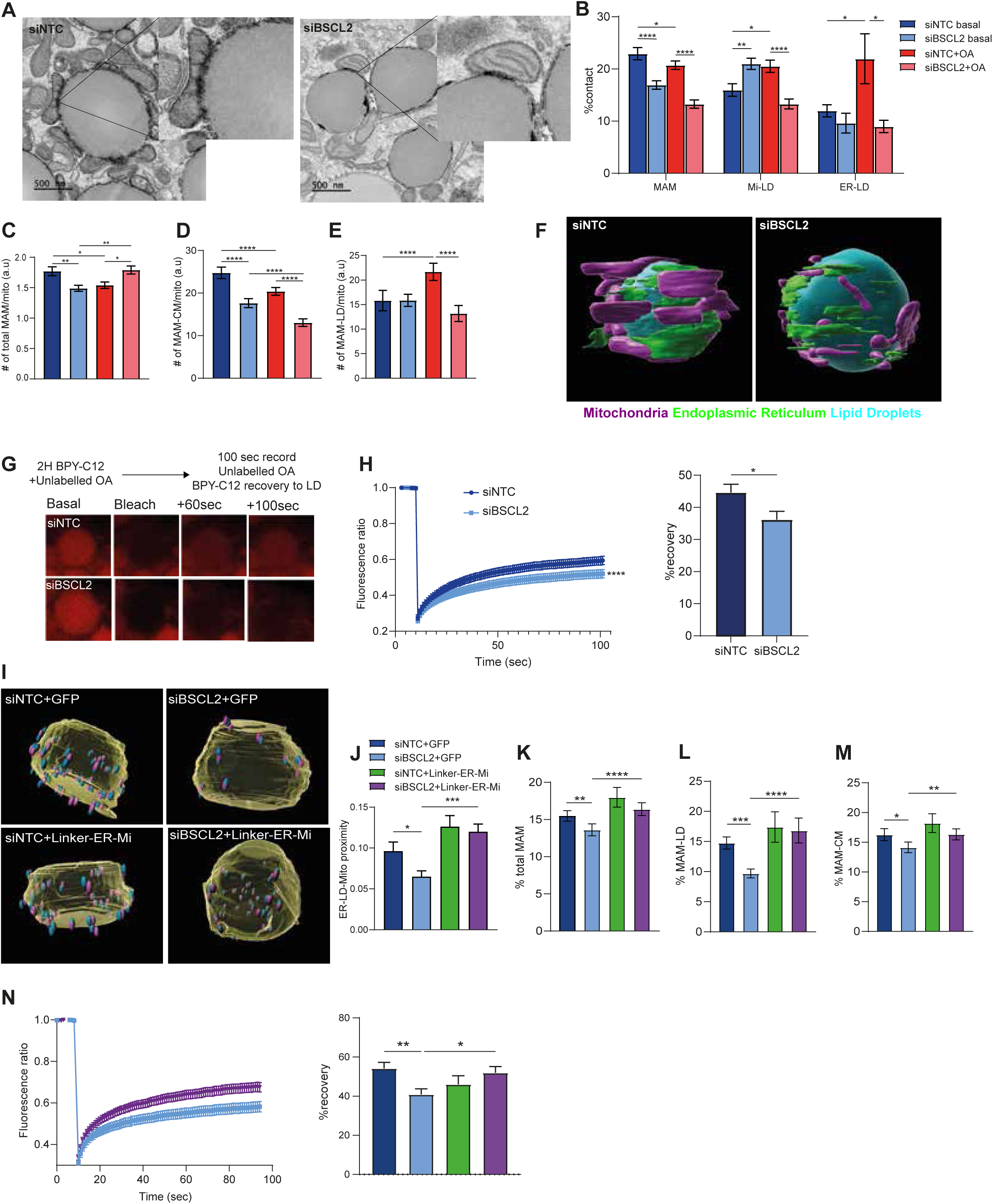
ER-to-LD lipid flux is altered is seipin absence, and restored by MAM-LD restructuration. 3T3-L1 adipocytes, after 7 days of differentiation, were reverse-transfected with the indicated siRNAs. After 72 hours, cells were incubated in low-glucose media ± oleic acid for 6 hours. TEM images (A) and analysis of quantified MCS (B), MAM number (C), and MAM surface area involving CM (D) or in contact with LDs (E). Representative images of FIB-SEM acquisitions on differentiated 3T3L1 with the indicated siRNA (F). Fluorescence recovery after photobleaching (FRAP) was performed in differentiated 3T3-L1 adipocytes (G-H). Kinetics of fluorescence recovery at LDs are shown, and percentage recovery is quantified. Bars: mean ± SEM, *n* > 120 cells/sample, 3 independent experiments. 3T3-L1 adipocytes were reverse-transfected with the indicated siRNAs and infected with adenovirus expressing GFP or Linker-ER-Mi. Structured Illumination Microscopy (SIM) was used to visualize mitochondria (Tom20), ER (PDI), and LDs (PLIN1), and spatial proximity was analysed with Imaris (I-J). Representative images show PLIN1 (yellow, segmented as LDs), PDI (blue, segmented as ER spots), and Tom20 (segmented mitochondria). TEM quantified the percentage of mitochondria engaged in MAMs (K), the proportion of MAMs contacting LDs (L), and those involving CM (M). FRAP analyses were performed (N): kinetics and percentage recovery are shown. Bars: mean ± SEM, *n* > 120 cells/sample, 3 experiments.

Next, we evaluated neutral lipid incorporation into LDs using fluorescence recovery after photobleaching (FRAP). Cells were incubated with the fluorescent fatty acid BODIPY-C12 (BPY-C12) for two hours, during which it is metabolized and esterified into neutral lipids, predominantly TG, that accumulate in LDs. After 2 hours, photobleaching was applied to LDs and fluorescence recovery was monitored. Seipin-deficient cells exhibited markedly reduced recovery, indicating impaired TG transfer (Figure 2G-H).

The above data are consistent with previous reports showing reduced ER-to-LD lipid flux in the absence of seipin ^13,14^. However, whether this defect depends on MAMs remains unclear. To test whether MAM–LD contacts directly mediate ER-to-LD lipid transfer, we genetically modulated contact formation by overexpressing Linker-ER-Mi, an artificial ER-mitochondria tether designed to promote MAM assembly^27^. We expressed the Linker-ER-Mi in seipin deficient adipocytes and we used Structured illumination microscopy (SIM) to assess the effect on MAM-LD. SIM analysis with immunolabeling for mitochondria (Tom20), ER (PDI), and LDs (PLIN1), followed by 3D analysis using a 380 nm proximity threshold^24^, showed that Linker-ER-Mi restored the spatial proximity between ER, mitochondria, and LDs (Figure 2I-J). TEM further confirmed that Linker-ER-Mi expression reestablished MAM structures, particularly MAM-LD contacts (Figure 2K-M). Repeating the FRAP assay under these conditions demonstrated that Linker-ER-Mi fully rescued the ER-to-LD lipid flux defect in siBSCL2-treated adipocytes (Figure 2N).

Altogether, these results show that seipin deficiency decreases MAM-LD contacts and ER-to-LD TG flux, and that artificially increasing MAM-LD contacts can restore lipid transfer even in the absence of seipin.

### Seipin regulates LD lipid storage by modulating MAM–LD contacts in a calcium-dependent manner

Given the known role of MAMs in ER-to-mitochondria calcium signaling, we next performed dynamic calcium imaging. In 3T3-L1 adipocytes, using Rhod-2 to monitor mitochondrial calcium uptake following IP3-induced ER calcium release, we found that seipin deficiency significantly reduced the calcium peak, and this defect was reversed by Linker-ER-Mi expression (Figure 3A). These findings raise the possibility that calcium exchange may contribute to the corrective effect of Linker-ER-Mi on ER-to-LD lipid transfer. To test whether the effect of Linker-ER-Mi on calcium transfer is required for its ability to rescue lipid flux in seipin-deficient cells, we blocked mitochondrial calcium uptake by silencing the mitochondrial calcium uniporter (MCU) using siRNA. We confirmed that siMCU effectively reduced MCU mRNA levels and abolished the Linker-ER-Mi-induced increase in mitochondrial calcium import (Supplemental 1A-B). We then performed FRAP experiments and found that inhibition of mitochondrial calcium uptake by MCU knockdown abolished the rescue of ER-to-LD lipid transfer normally observed with Linker-ER-Mi (Figure 3B-C). Importantly, MCU knockdown did not alter the structural proximity between the three organelles (ER, mitochondria, and LDs), as shown in Figure 3D-E, highlighting the critical role of calcium in regulating ER-to-LD lipid flux. Given the importance of calcium in mitochondrial function—particularly in the activation of enzymes in the TCA cycle—we next assessed the phosphorylation state of pyruvate dehydrogenase (PDH), a key enzyme that converts pyruvate into citrate. While PDH phosphorylation, which inhibits its activity, is promoted under low-calcium conditions, we observed an elevation of PDH phosphorylation levels in seipin-deficient adipocytes that returned to baseline upon Linker-ER-Mi expression (Figure 3F-G). Of note, PDH phosphorylation levels were also elevated in the AT of iATSKO mice, further supporting that mitochondrial calcium import impairment is a genuine hallmark of seipin deficiency (Figure 3H-I). Finally, by tracing ^13^C-labeled glucose metabolism, we found that citrate production was impaired in seipin-deficient cells and that this defect was corrected by Linker-ER-Mi expression (Figure 3J). Altogether, our findings indicate that calcium flux from the ER plays a crucial role in mediating triglyceride transfer from the ER to LD.

**Figure 3:**
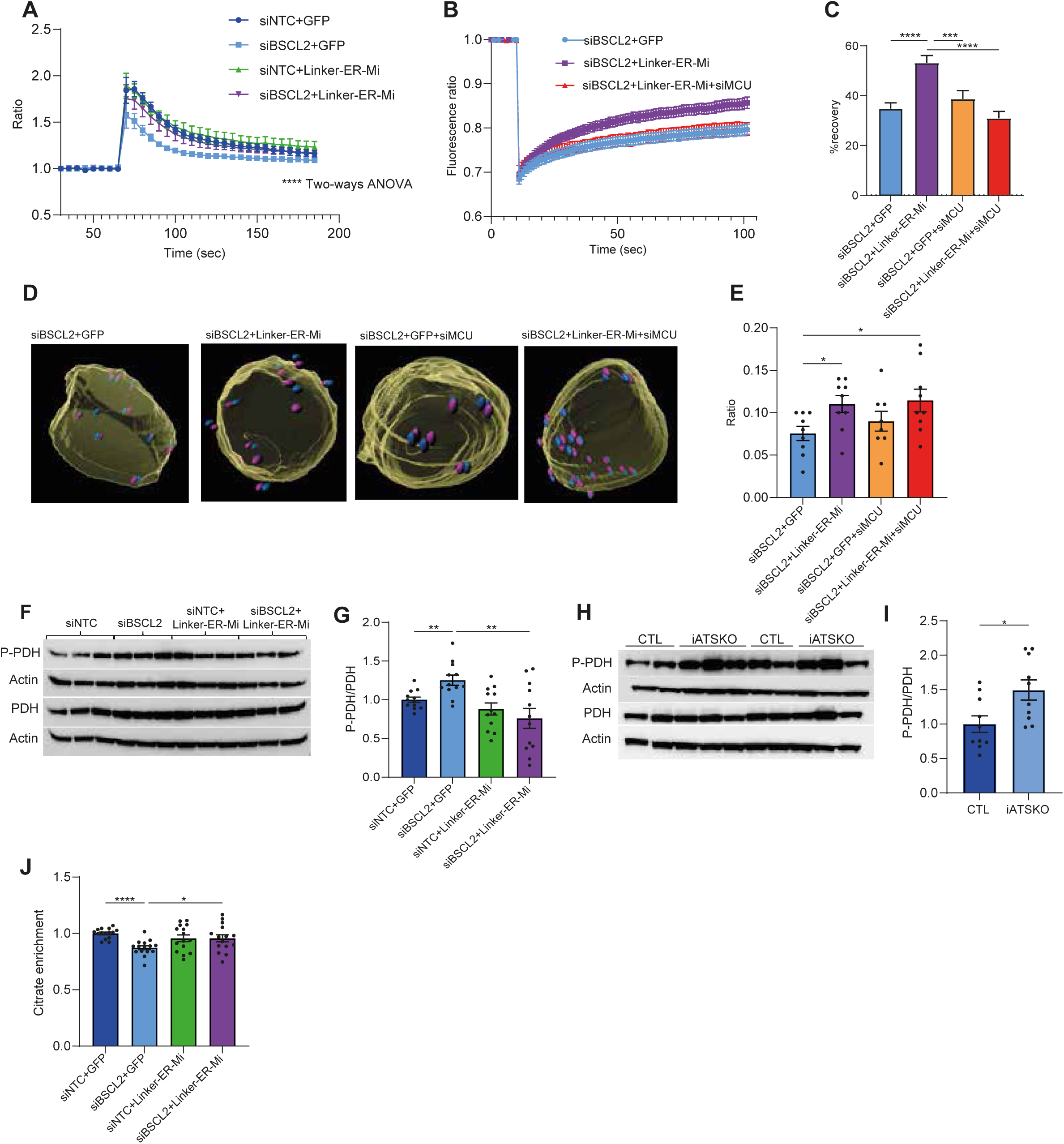
MAM-LD control lipid flux in a calcium dependant-way. 3T3-L1 adipocytes were reverse-transfected with the indicated siRNAs and infected with adenovirus expressing GFP or Linker-ER-Mi. Mitochondria calcium was monitored using caged IP₃ and Rhod2. After 60 seconds of basal acquisition, UV flash photolysis was applied to release IP₃. The Y-axis represents the fluorescence ratio over basal (A). FRAP analysis was performed in differentiated 3T3-L1 cells (B-C), showing fluorescence kinetics (B) and recovery percentage (C). SIM imaging was used to assess mitochondria (Tom20), ER (PDI), and LDs (PLIN1), and spatial analysis was performed with Imaris (D-E). Representative segmented images are shown (right). Western blot images (F) and quantification (G) were performed on 3T3-L1 adipocytes 4 days post-siRNA transfection. Western blot images (H) and quantification (I) were performed on inguinal adipose tissue lysates from control and iATSKO mice. Bars represent mean ± SEM, n = 8 per genotype; samples were run twice in WB. *P* < 0.005, Mann–Whitney test. Fractional enrichment of [¹³C]-glucose–derived citrate was analyzed by LC-HRMS (J). Bars: mean ± SEM, each condition in triplicate, *n* = 3 experiments. ***p*** < 0.005, Mann–Whitney test.

### Mapping MCS during adipogenesis and under distinct nutritional conditions

Our work on seipin deficient adipocytes demonstrates that alterations in MCS involving the ER, LDs, and mitochondria and especially the tripartite MAM-LD, are associated with severe adipocyte dysfunction. As those contact sites have been poorly studied in adipocytes, we characterized these interactions during two key physiological situations of adipocyte function: adipogenesis and the response to nutrient availability in differentiated cells.

To assess MCS during adipogenesis in 3T3-L1 cells, we performed transmission electron microscopy (TEM), which revealed that the appearance of LDs at day 4 post-induction of differentiation was accompanied by a redistribution of mitochondria. From this stage onward, the images highlighted the presence of peridroplet mitochondria (PDM) surrounding LDs. Quantification of the MCSs showed an increase in Mi–LD contacts and a concomitant decrease in Total-MAMs (Figure 4 A-B). To further decipher the specific implication of MAM-LD in this process, we segregated the quantification of MAM-CM, MAM-LD and Mi-LD. We confirmed a decrease of MAM-CM during adipocyte differentiation while MAM-LD were increased at day 4 of induction, highlighting specific regulation of this subtype of MAMs during adipocyte differentiation (Figure 4C).

**Figure 4:**
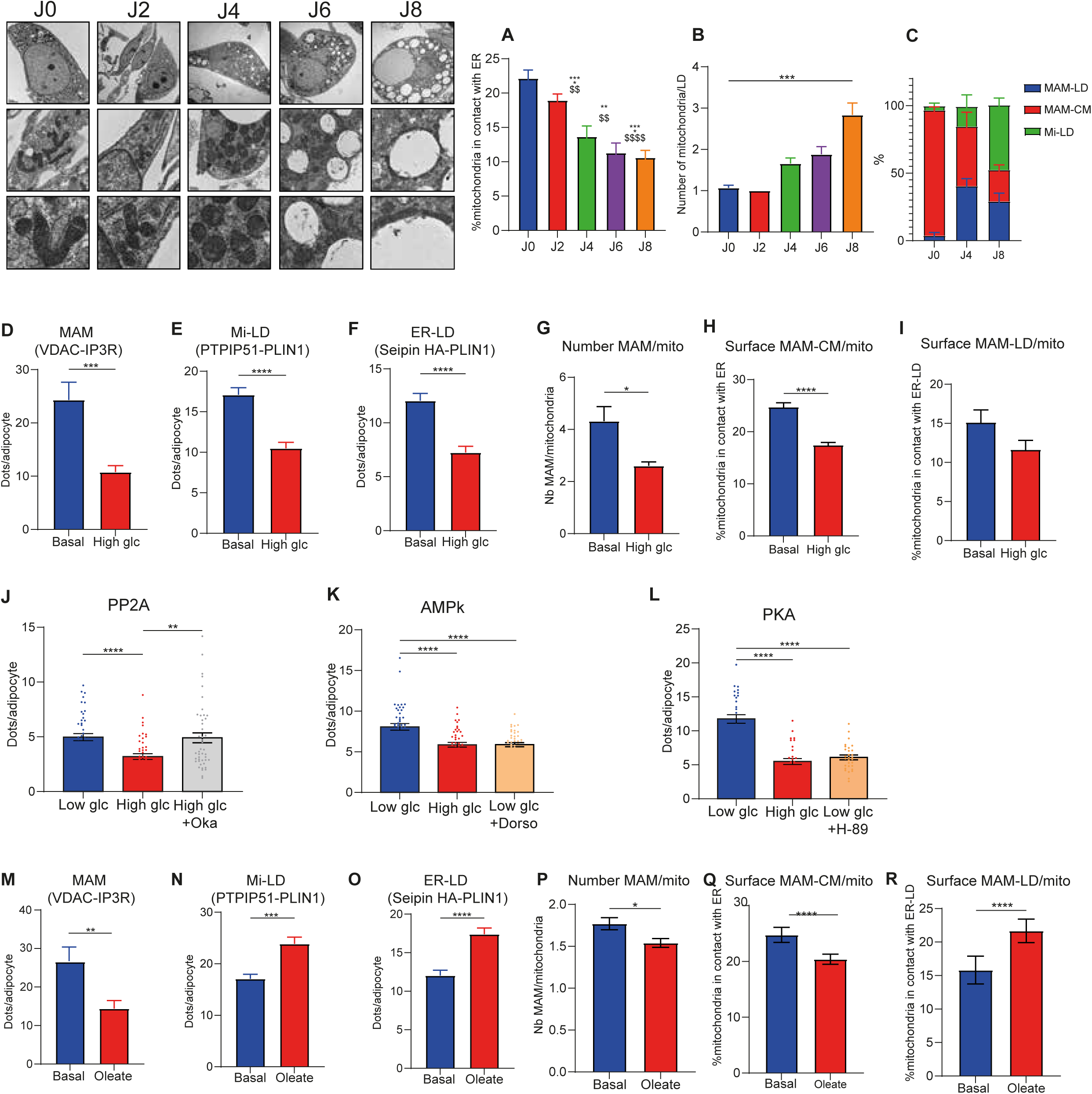
Mapping MCS during adipogenesis and under distinct nutritional conditions. 3T3-L1 differentiation is initiated at day 0 with the addition of the adipogenic cocktail (see method section). TEM is performed at the indicated day after differentiation induction. The percentage of mitochondria in contact with the ER (A), the number of mitochondria/LD contacts (B), and the different MAMs subtype (C). 3T3-L1 adipocytes after 7 days of differentiation were plated in 8-wells IBIDI. Cells were incubated in low-glucose (5mM) media +/- oleic acid (700µM), or +/- high glucose (25mM) for 6 hours. PLAs were performed to detect: ER-mitochondria contacts (VDAC/IP3R; D,M), mitochondria-LD contacts (PTPIP51/PLIN1; E,N) and ER-LD contacts (Seipin HA-PLIN1; F-O). Cells were incubated in low/high-glucose media and treated +/- okadaic acid (OA 10nM), Dorsomorphin (Dorso, 5µM), H-89 (10µM). PLA was performed to detect ER-mitochondria contacts (VDAC/IP3R; J,K,L). Bars: mean ± SEM, *n* > 120 cells/sample, 3 experiments. The number of MAM per mitochondria (G,P), and the surface of MAM-CM (H,Q) or MAM-LD (I,R) were also analyzed by TEM using Fiji. Bar: mean +/- SEM, n>5 cells/group. *p<0.05, ***p<0.001, ****p<0.0001, Mann-Whitney test.

Then, we intended to decipher the MCS remodeling in response to nutrient treatment. Using PLA, we quantified MAMs through IP3R–VDAC interactions, Mi–LD contacts via PTPIP51–PLIN1, and ER–LD contacts via Seipin-HA–PLIN1. High glucose (25mM) exposure reduced all three types of contact sites (Figure 4D-F). TEM analysis revealed that only the decrease in MAM-CM reached statistical significance, whereas the reduction in MAM-LD did not (Figure 4G-I). The glucose-induced decrease in MAMs was abolished by treatment with okadaic acid (Oka), an inhibitor of protein phosphatase 2A (PP2A), which is known to be activated by xylulose-5-phosphate, a product of glucose metabolism (Figure 4J). Conversely, the increase in MAMs observed under low-glucose (5mM) conditions was blunted by dorsomorphin and H89, inhibitors of AMPK and PKA, respectively—two signalling pathways typically activated in response to nutrient deprivation (Figure 4K-L). Interestingly, in fully differentiated 3T3-L1 adipocytes, treatment with oleic acid (OA) led to a reduction in MAMs, while ER–LD and Mi–LD contact sites were significantly increased (Figure 4 M-O). However, while lipid loading decreased MAM-CM, it promoted MAM-LD formation (Figure 4P-R). Thus, glucose and lipids have different effects on MCS remodeling in adipocytes.

Together, these data strongly suggest that MCS remodelling is a key determinant of adipocytes metabolic flexibility and that MAM-LD plays a key role in lipid handling.

### MCS Nutritional Remodelling Is Abolished in the Adipose Tissue of Diet Induced Obese Mice

Seipin deficiency is a severe and primary state of adipocyte dysfunction. We next ask whether MCS abnormalities are also a hallmark of the most common form of obesity-related adipocyte dysfunction in mice. Obesity was induced by high-fat diet (HFD) feeding for 3 months, resulting in significant weight gain and systemic insulin resistance (Supplemental Fig. 4A-B). Histological analysis confirmed adipocyte hypertrophy in the gonadal white AT (gWAT) of obese mice compared to control lean mice (Supplemental 1F). Using PLA, we found that in lean mice, feeding decreased MAMs and increased contacts involving LDs, including ER–LD and Mi–LD contacts. In contrast, these adaptive changes were absent in obese mice: fasting did not increase MAMs, and the postprandial rise in ER–LD and Mi–LD contacts was also missing (Figure 5A–C).

**Figure 5:**
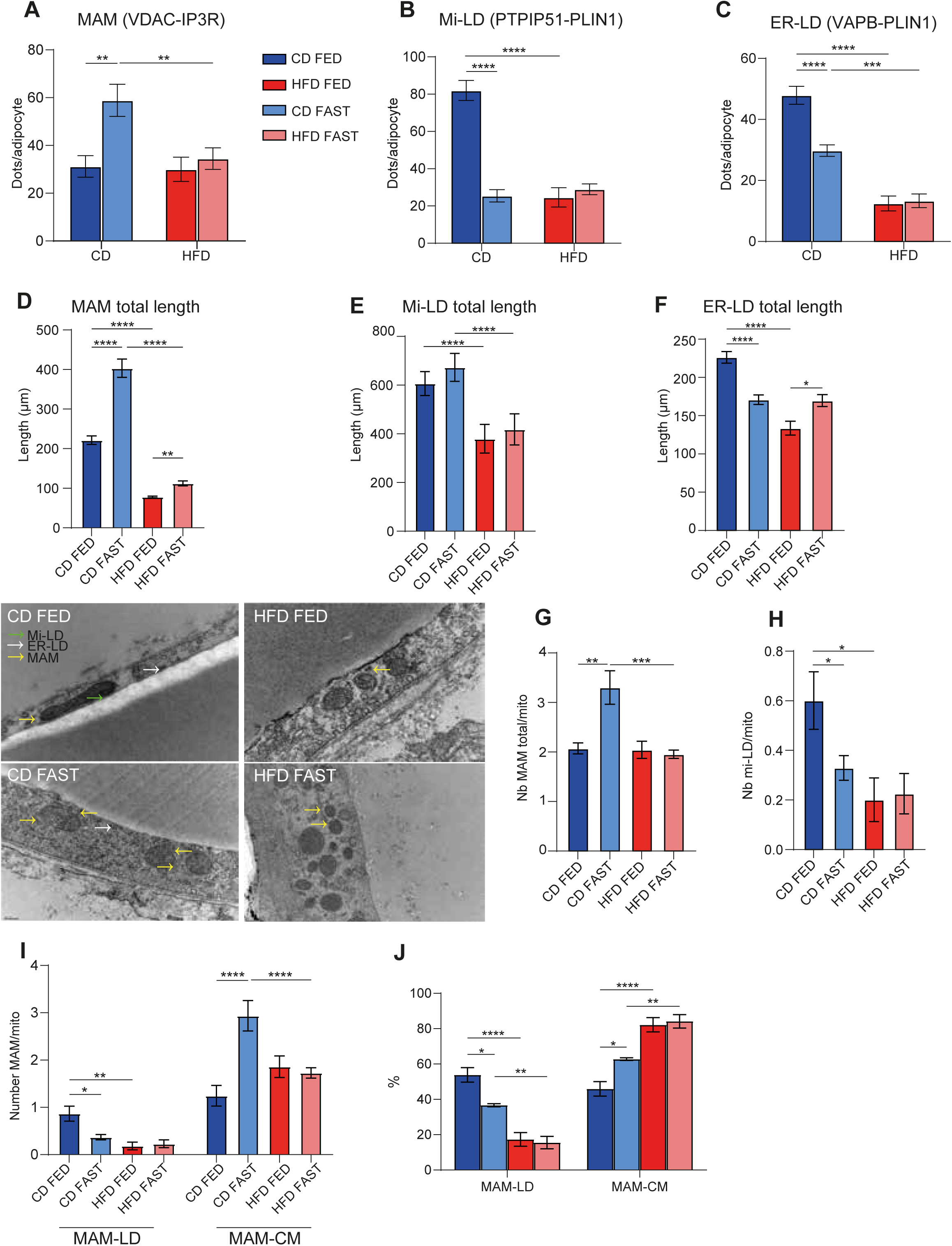
MCS are altered in AT of obese mice. Cryopreserved gonadal adipose tissue from mice fed chow diet or high fat diet for 3 months was collected in fed and 18-hour fasted state. PLAs quantified the following contacts: ER-mitochondria (VDAC/IP3R, A), mitochondria-LD contacts (PTPIP51/PLIN1, B), and ER-LD contacts (VAPB-PLIN1, C). Bars: mean +/- SEM, n=5 mice per group, >100 cells per mouse. *p<0.05, ***p<0.001, ****p<0.0001, Mann-Whitney test. Gonadal adipose tissue from mice fed chow diet or high fat diet for 3 months were collected in fed or 18-hour fasted state. Transmission electron microscopy was used to quantify the following organelle contacts length with FIJI : ER-mitochondria contact (MAM, D), mitochondria-LD contact (Mi-LD, E), and ER-LD (F). TEM was also used to quantify the number of total MAM (G) and mi-LD contact per mitochondria (H). Number (I) and proportion (J) of MAM-LD and MAM-CM were further distinguished. Bars: mean +/- SEM, n=3 mice per group, 5 cells per mouse. *p<0.05, ***p<0.001, ****p<0.0001, Mann-Whitney test

TEM analysis in control mice showed that fasting increased both the length and number of MAMs (Figure 5D, G), while feeding increased ER–LD contact length (Figure 5F). Mi–LD contact length was not affected by fasting (Figure 5E); however, the number of Mi–LD contacts decreased upon fasting, as quantified by PLA (Figure 5H). When normalized to organelle surface area, Mi–LD contact surface was reduced in the fed state, whereas ER–LD contact area remained stable (Supplemental 1C–E). In obese mice, all three MCS types were reduced during fasting. Furthermore, the expected postprandial increases in Mi–LD and ER–LD contacts were absent, and the MAM remodelling response was strongly blunted.

Feeding exerted opposite effects on the two MAM subtypes in lean mice: MAM–LDs increased, while MAM–CMs decreased. In obese mice, the fasting-induced increase in MAM–CMs was abolished, and MAM–LD numbers did not rise after feeding, indicating a selective disruption in the remodelling of tripartite contact sites (Figure 5I–J).

Altogether, these findings demonstrate that systemic insulin resistance and obesity are associated with a marked impairment in MCS remodelling in gonadal AT, highlighting a loss of metabolic flexibility at the level of organelle interactions.

### MCS disruption is a causal and reversible trigger of adipocyte dysfunction

As in vivo, MCS abnormalities is associated with AT dysfunction and systemic metabolic inflexibility, we used FATE1 ectopic expression in adipocytes to assess the consequences of MAMs disruption. FATE1 is normally expressed in the testis but not in adipocytes, where it acts as a spacer between the ER and mitochondria, thereby decreasing MAM. We overexpressed FATE1 in adipocytes and assessed MAM and other MCS using PLA. FATE1 expression significantly reduced both MAM and ER-LD contact sites under basal and oleate-treated conditions, while mi-LD increased in the basal state only (Figure 6A-C). To assess whether MAM disruption impairs metabolic flexibility, we measured glycerol release following lipolytic stimulation by the β-adrenergic agonist CL-316,243. In control cells, glycerol release was robustly induced (Figure 6D) while this response was significantly blunted in FATE1-expressing cells. Furthermore, FATE1 expression decreased insulin-stimulated Akt phosphorylation by 30%, indicating reduced insulin sensitivity (Figure 6E). Given the strong alteration of MCS remodeling in the lipid loading state, we measured ER-to-LD lipid flux by using FRAP. FATE1 overexpression decreased fluorescence recovery, highlighting lipid storage impairment (Figure 6F). In order to better characterize MCS organization in this model, we quantified MAM, MAM-LD and MAM-CM using PLA MAM (VDAC-IP3R), combined with IF for PLIN1 imaging withSIM. Total MAM were strongly decreased in FATE1 overexpressing-adipocytes, and this decrease is mainly explained by the decrease of MAMs-LD, as MAMs-CM are not significantly reduced (Figure 6 G-I)

**Figure 6:**
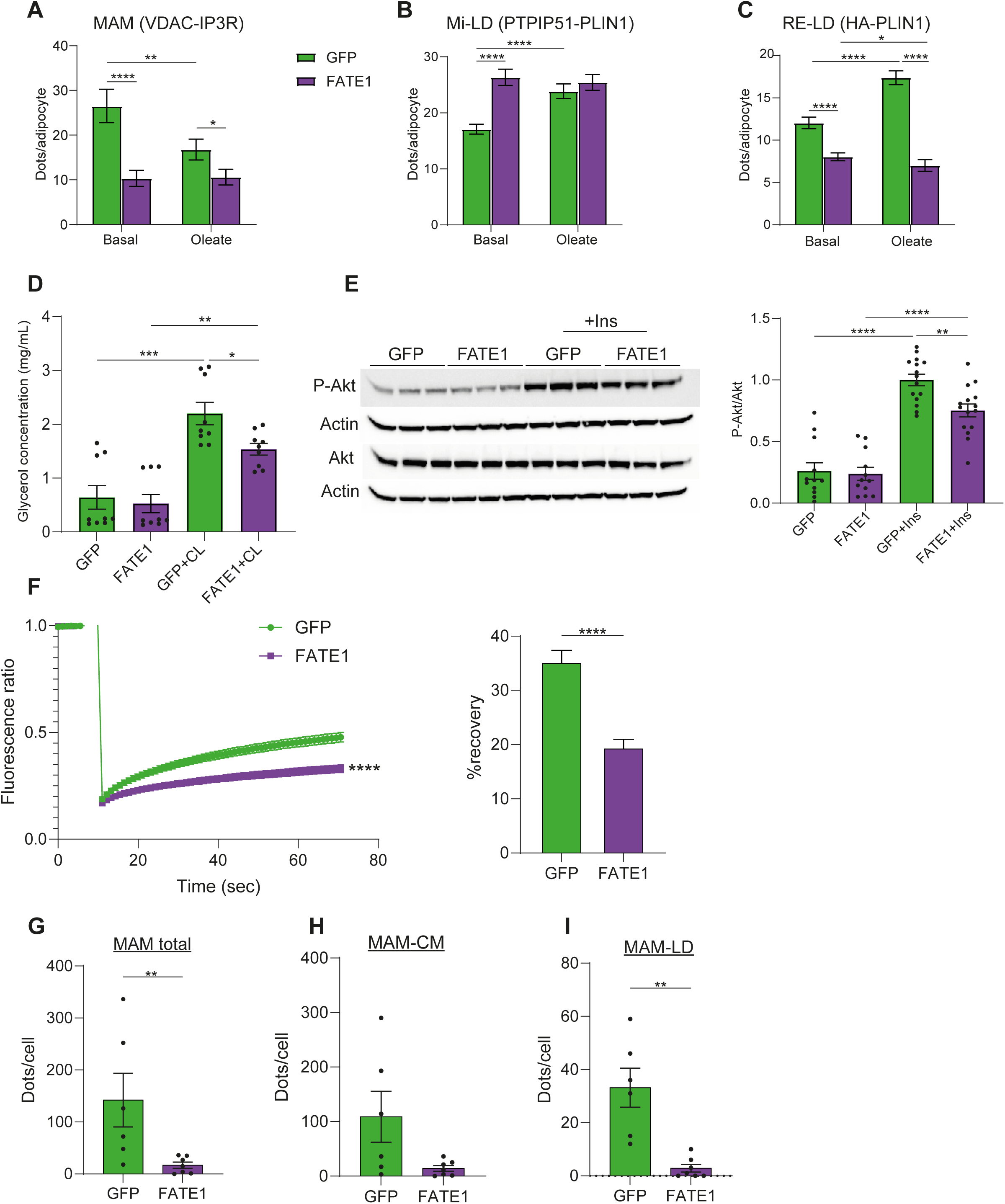
MAM disruption alters metabolic flexibility and impairs lipid storage. 3T3-L1 adipocytes, after 7 days of differentiation were infected with adenovirus expressing GFP or FATE1. Cells were incubated in low-glucose media +/- oleic acid for 6 hours. PLAs were performed to detect: ER-mitochondria contacts (VDAC/IP3R; A), mitochondria-LD contacts (PTPIP51/PLIN1; B) and ER-LD contacts (Seipin HA-PLIN1; C). Lipolysis was measured by quantifying glycerol release in extracellular media after 2 hours of incubation +/- CL316-243 (D). Akt phosphorylation was assessed by Western Blot and quantified (E). FRAP analysis was performed (F) fluorescence kinetics (left) and recovery percentage are shown (right). Bars: mean +/- SEM, each condition in triplicate, n=3 experiments, *p<0.05, ***p<0.001, ****p<0.0001, Mann-Whitney test. The number of mitochondria-associated membranes (MAMs) was quantified using a proximity ligation assay (PLA) for IP3R and VDAC, combined with PLIN1 immunofluorescence, and analyzed by Structured Illumination Microscopy (G-I). Total PLA dots were analyzed (G), and further distinguished by their distance from lipid droplets (LD): MAM-CM (>380 nm from LD; H) and MAM-LD (<380 nm from LD; I) Bars: mean +/-SEM, each condition in duplicate, n=2 experiments, *p<0.05, ***p<0.001, ****p<0.0001, Mann-Whitney test.

In order to get insights more specifically in the role of MAM-LD in adipocyte physiology, we disrupted MCS organization by deleting PTPIP51, a mitochondrial tether protein localized to both MAMs and Mi–LD contact sites. Using CRISPR/Cas9, we generated PTPIP51 knockout (KO) 3T3-L1 cells. Knockout efficiency was validated by Western Blot (Supplemental 1A), and both populations retained the ability to differentiate into adipocytes (Supplemental 1B). PLA analysis revealed that oleic acid failed to induce the expected MCS remodelling in PTPIP51 KO cells (Figure 7A-C). Calcium signalling and oxygen consumption were altered in KO cells (Supplemental 1C-D). Lipolysis assays revealed a ∼40% decrease in both basal and CL-316,243-induced glycerol release in PTPIP51 KO cells (Figure 7D). Insulin-induced Akt phosphorylation was also reduced by approximately 50% in KO cells compared to controls (Figure 7E). To ensure the specificity of our model, we overexpressed PTPIP51 in KO PTPIP51 cells and observed a rescue of both calcium flux and the lipolysis defect (Supplemental 1 E-F).

**Figure 7:**
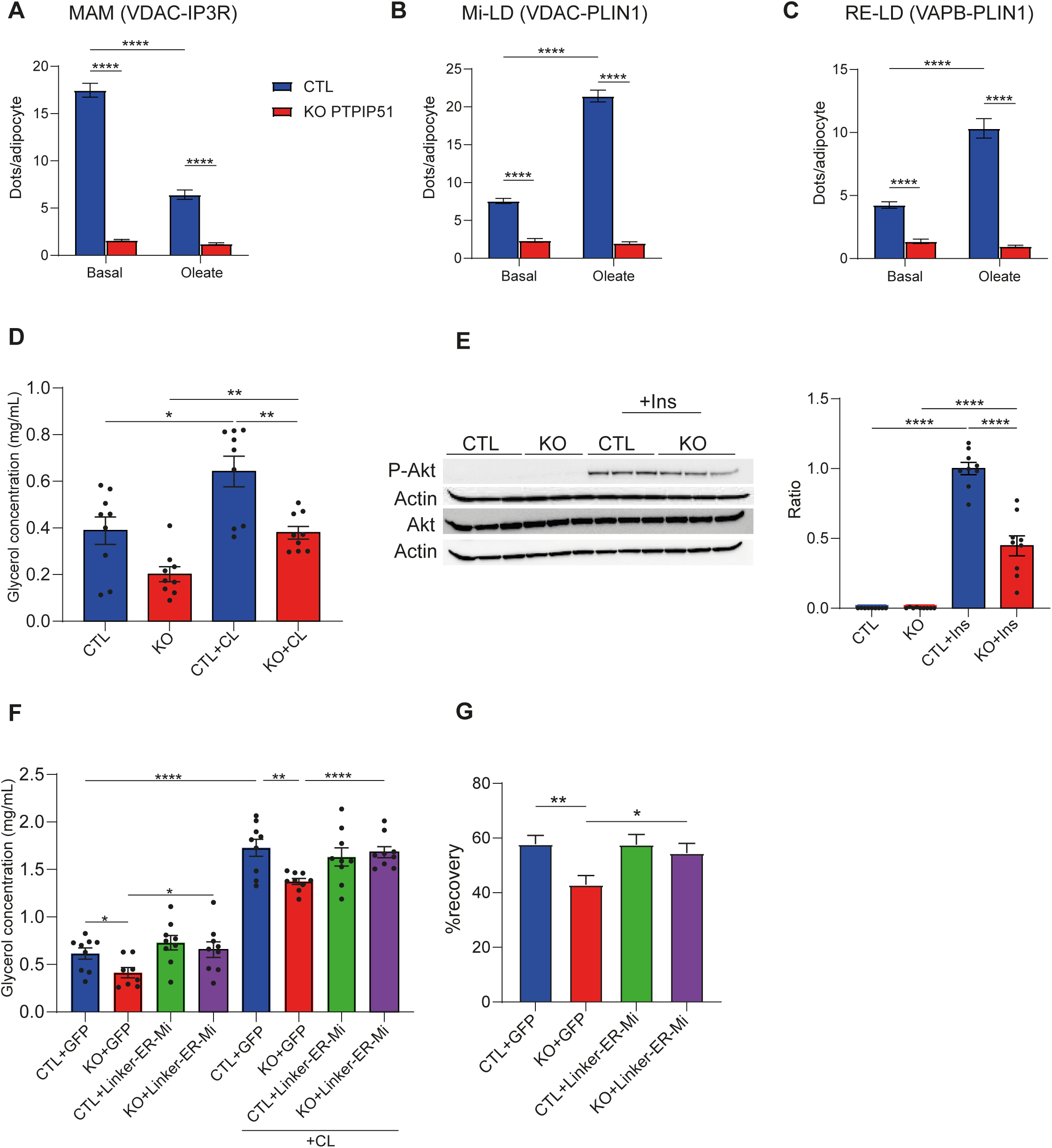
MAM reinforcement in MCS altered cellular model restores adipocyte properties. CTL and KO PTPIP51 3T3-L1 adipocytes, were incubated in low-glucose media +/- oleic acid for 6 hours. PLAs were performed to detect: ER-mitochondria contacts (VDAC/IP3R; A), mitochondria-LD contacts (VDAC/PLIN1; B) and ER-LD contacts (VAPB-PLIN1; C). Lipolysis was measured by quantifying glycerol release in extracellular media after 2 hours of incubation +/- CL316-243 (D). Akt phosphorylation was assessed by Western Blot and quantified (E). Bars: mean +/- SEM, each condition in triplicate, n=3 experiments, *p<0.05, ***p<0.001, ****p<0.0001, Mann-Whitney test. CTL and KO PTPIP51 3T3-L1 adipocytes were infected with adenovirus expressing GFP or Linker-ER-Mi.). Lipolysis was measured by quantifying glycerol release in extracellular media after 2 hours of incubation +/- CL316-243 (F). FRAP analysis was performed and recovery percentage was calculated (G). Bars: mean +/- SEM, each condition in triplicate, n=3 experiments, *p<0.05, ***p<0.001, ****p<0.0001, Mann-Whitney test.

Finally, to determine whether the functional defects observed in *PTPIP51*-deficient adipocytes were specifically due to impaired MAM-LD organization, we expressed Linker-ER-Mi to artificially reinforce MAMs. To assess the functional relevance of MAM reinforcement, we performed lipolysis assays. In *PTPIP51* KO cells, lipolytic capacity was impaired, but was fully rescued upon Linker-ER-Mi expression (Fig. 7F). Using FRAP analysis, we confirmed that ER-to-LD lipid transfer was reduced in *PTPIP51* KO adipocytes. Strikingly, expression of Linker-ER-Mi fully restored fluorescence recovery kinetics, indicating a rescue of ER-to-LD lipid flux (Figure 7G). This suggests that strengthening ER–mitochondria contacts, thus restoring MAM-LD architecture, is sufficient to restore key adipocyte functions disrupted by *PTPIP51* loss.

## Discussion

Understanding how metabolic flexibility is regulated in adipocytes is crucial to pave the way for new therapeutic strategies to tackle the metabolic complication associated with obesity. Here, starting with the study of an extreme and rare case of primary adipocyte dysfunction, the CGL due to seipin deficiency, we studied the role of MCS involving the ER, the mitochondria, and the LD in adipocytes homeostasis. Indeed, we highlight that the MAM-LD tripartite contact sites are promoted by lipid loading and that they regulate lipid flux from the ER to the LD in a calcium-dependent manner. Then, we established the landscape of MCS remodelling under nutritional stimulation and demonstrated that those adaptations are lost in the AT of obese mice. We further demonstrate that specific MCS disturbances are sufficient to induce lipolysis and insulin signalling alteration in adipocytes. Interestingly, the abnormalities associated with MCS disruption were corrected by the use of a synthetic linker, the Linker-ER-Mi, that reinforces MAMs. Altogether, we highlight the key role of MAM-LD in lipid handling and the importance of MCS involving the ER, the LD and the mitochondria, in metabolic flexibility.

Seipin is a critical regulator of adipocyte homeostasis, since its deficiency leading to severe adipocyte dysfunction and lipoatrophy in both humans and mice. Seipin is known to be enriched at endoplasmic reticulum–lipid droplet (ER–LD) contact sites, and in fibroblasts derived from patients with seipin deficiency, these contacts appear disorganized^14^. Additionally, seipin is enriched at MAMs, and its deficiency disrupts calcium homeostasis^15,24,28^. In this study, we demonstrate that seipin deficiency reduces all three contact types involving the ER, mitochondria, and LD, with a particularly pronounced effect on MAM-LD contacts. Lipidomic analysis of LDs isolated from iATSKO mouse AT revealed decreased TG and increased diacylglycerol DG content, suggesting a defect in TG synthesis or transport. Using FRAP analysis, we showed that seipin deficiency impairs ER-to-LD lipid transfer *in vitro*, and that reinforcement of MAM-LD contacts can restore this lipid transfer. Although we lack specific tools to selectively reinforce MAM-LD contacts, we employed Linker-ER-Mi, which increases proximity between the ER and mitochondria. Using both SIM and TEM, we confirmed that, in seipin-deficient adipocytes, Linker-ER-Mi primarily reinforces MAM-LD contacts. To further decipher the mechanism of action, we knocked down the MCU in order to prevent the calcium entry into mitochondria. This intervention abolished the effect of Linker-ER-Mi, demonstrating that the restoration of lipid flux is calcium-dependent. Thus, beyond restoring the structural integrity of MAM-LD, recovering the calcium exchange is essential. Supporting this, studies in hepatocytes from yellow catfish have shown that phosphatidic acid loading promotes lipid deposition by recruiting seipin to MAMs^29^. In their model, seipin deficiency did not alter MAM structure or number, but did prevent lipid accumulation in a calcium-dependent manner, further highlighting the crucial role of calcium signalling modulation in the effect of MAMs on lipid storage. The precise mechanisms by which calcium regulates lipid storage remain to be elucidated. It is plausible that calcium controls ATP production required for lipid transfer protein function, that some of these proteins are directly calcium-dependent, or that calcium regulates citrate production, the initial substrate for lipogenesis. Notably, we observed that citrate production is decreased in seipin-deficient adipocytes and restored by Linker-ER-Mi expression.

How can we integrate our findings with previous research on seipin function? A substantial body of work demonstrates that seipin forms oligomeric, donut-like complexes whose central hydrophobic helices bind and concentrate triacylglycerols (TAG), thereby facilitating LD formation^30–33^. Loss of seipin disrupts ER membrane curvature, and seipin preferentially localizes to tubular ER subdomains rather than sheet-like regions^34^. Since the reinforcement of MAM-LD contacts is sufficient to restore lipid flux even in the absence of seipin, we propose that seipin plays a central role in organizing MAM-LD contacts during lipid loading, thereby enabling efficient TG synthesis and storage in the LD. Notably, a recent study that thoroughly analyzed the protein composition of MAM-LD contacts demonstrated that these sites are enriched in enzymes involved in TG synthesis, such as GPAT, AGPAT2, LIPIN, and DGAT2^23^. Strikingly, seipin has been shown to interact with GPAT3^35^, LIPIN^36^ and AGPAT2^37^. However, our lipidomic analysis did not reveal differential enrichment of these proteins in the LD environment of seipin-deficient AT compared to controls. Therefore, we do not propose that seipin directly controls their recruitment. Instead, we suggest that seipin plays a central role in scaffolding the MAM-LD contacts, determining their structural properties to ensure their functional role in LD lipid storage.MAM-LD contacts have only recently been named and characterized^23,24^, and we discuss our findings in the context of emerging literature to better understand their role in lipid handling. Using TEM and PLA in both AT and 3T3-L1 adipocytes, we show that lipid loading increases contacts involving the LD, specifically ER/LD and mitochondria/LD (Mi/LD) contacts. However, lipid loading exerts opposite effects on MAM subtypes: oleic acid increases MAM-LD contacts while decreasing MAM-CM contacts. Consistently, in the liver, oleic acid has been reported to decrease MAM-CM and increase MAM-LD contacts^23^. Therefore, we are proposing that the MAMs involving CM would be the “classical” MAMs, known to support catabolic processes during fasting^17^, while those involving PDM are requested during lipid loading challenge. Beyond seipin deficiency, we demonstrate that disruption of MAM-LD contacts by FATE1 overexpression, as well as global MCS defects in PTPIP51 knockout adipocytes, severely impairs lipid transfer from the ER to the LD. As observed in seipin-deficient adipocytes, Linker-ER-Mi rescues this defect in PTPIP51-deficient cells, supporting the hypothesis that the effect of PTPIP51KO relies indeed on its action in structuring MCS^38^. In HeLa cells, the lipid transfer proteins ORP5 and ORP8 orchestrate LD biogenesis at MAM-LD sites by interacting with seipin and controlling its localization ^24^. In adipocytes, MAM-LD contacts have not yet been directly explored. One study reported that the MAM-localized mitochondrial fission protein DRP1 is involved in lipid trafficking to LDs, suggesting a need for cooperation among all three organelles for LD biogenesis, though without directly demonstrating the involvement of MAM-LD ^39^. Additionally, the outer mitochondrial membrane protein MIGA2 reinforces mitochondria/LD contacts and promotes lipogenesis ^40^. We attempted to isolate MAM-LD structures, but our yields were insufficient for unbiased analysis, particularly in comparison to MAM-CM. Further studies are needed to precisely identify the key players and mechanisms that regulate lipid metabolism at MAM-LD contacts.

We evaluated the overall contribution of MCS to adipocyte metabolic flexibility. Glucose loading decreased MAM abundance, consistent with previous findings in the liver and supporting the idea that MAM formation are induced by fasting ^17^. We found that MAM regulation by glucose depends on AMPK and PKA, two key signaling pathways involved in adaptation to fasting. Similar findings were reported in the liver previously^41^. Lipolysis, which is stimulated by fasting and the PKA pathway^42^, was impaired by specific MAM depletion but restored by Linker-ER-Mi–mediated MAM reinforcement. Furthermore, disruption of MAMs induced cell-autonomous insulin resistance in adipocytes, reinforcing the role of MAMs as metabolic hubs essential for the regulation of metabolic flexibility.

Finally, having shown that MAM-LD alterations are associated with severe adipocyte dysfunction and that MCS remodeling is a key player in metabolic flexibility, we investigated whether MCS involving the ER, LD, and mitochondria are altered in the AT of obese mice—a model characterized by metabolic inflexibility. We demonstrated that nutrient-induced MCS remodeling is abolished in the AT of obese mice at a stage when they develop insulin resistance. Most precisely, while MAM-CM were increased in the AT of obese mice, MAM-LD were nearly absent. This urge us to better define what type of MAM are we talking about when we are describing the MCS. The state of MAM in the liver of obese mice has been largely debated. While some groups found them decreased ^43^ other found them increased ^44,45^. It would be interesting to further define what type of MAM is being discussed in these two reports. The combination of PLA and SIM analysis used here, based on a previous publication ^25^ might be an easy tools to determine MAM-LD state. Dysfunctional adipocytes in this context exhibit profound alterations in lipid and glucose metabolism, hallmarks of metabolic inflexibility ^46^. Particularly, reduced lipogenesis activity in the AT is a hallmark of insulin resistance and AT failure in the context of obesity ^47^. Based on our findings, we propose that abnormalities in MCS may contribute to this state of metabolic inflexibility, as in vitro, MCS disruption alone induces both insulin resistance and impaired lipolysis.

## Conclusion

Altogether, our work establishes MCS disruption as a hallmark of adipocyte dysfunction in two murine models of AT failure: CGL and diet-induced obesity. Specifically, contacts involving the ER, LDs, and mitochondria, are significantly impaired. Using synthetic linkers to reinforce these connections corrected several adipocyte abnormalities, validating MAM-LD as a potential therapeutic target. Given that MCS remodeling is crucial for metabolic flexibility, interventions like Linker-ER-Mi that clamp MAMs in a certain state, appear unpromising. Conversely, strategies that restore remodeling capacity or enable transient MAM-LD augmentation during the post-prandial phase represent promising future therapeutic avenues. This approach is particularly relevant considering the persistence of AT dysfunction (“fat memory”) even after weight loss. Targeting adipocyte dysfunction directly, alongside next-generation weight-loss medications, offers a synergistic therapeutic strategy.

## Supporting information

FIB SEM videos

## Acknowledgements

This study has been granted from Société francophone du diabète, Fondation Genavie, Société française d’endocrinologie, the FFRD (sponsored by Fédération Française des Diabétiques, Abbott, AstraZeneca, Eli Lilly, Merck Sharp & Dohme et Novo Nordisk) and the French National Research Agency (ANR-21-CE14-0024 MAMA and ANR-25-CE14-0269-01MICADO). We acknowledge the MicroPICell facility, SFR-Santé, INSERM, CNRS, UNIV Nantes, CHU Nantes, Nantes, France, member of the national infrastructure France-BioImaging supported by the French National Research Agency (ANR-10-INBS-04). We are most grateful to the Genomics and Bioinformatics Core Facility of Nantes (GenoBiRD, Biogenouest, IFB) for its technical support. We thank the SCIAM imagery platform, especially F.Manero for electron microscopy imaging and technical assistance. The present work has benefited from Imagerie-Gif core facility supported by l’Agence Nationale de la Recherche (FBI ANR-24-INBS-0005 (BIOGEN); SPS ANR-17-EUR-0007, EUR SPS-GSR)

---

**Supplemental 1.**
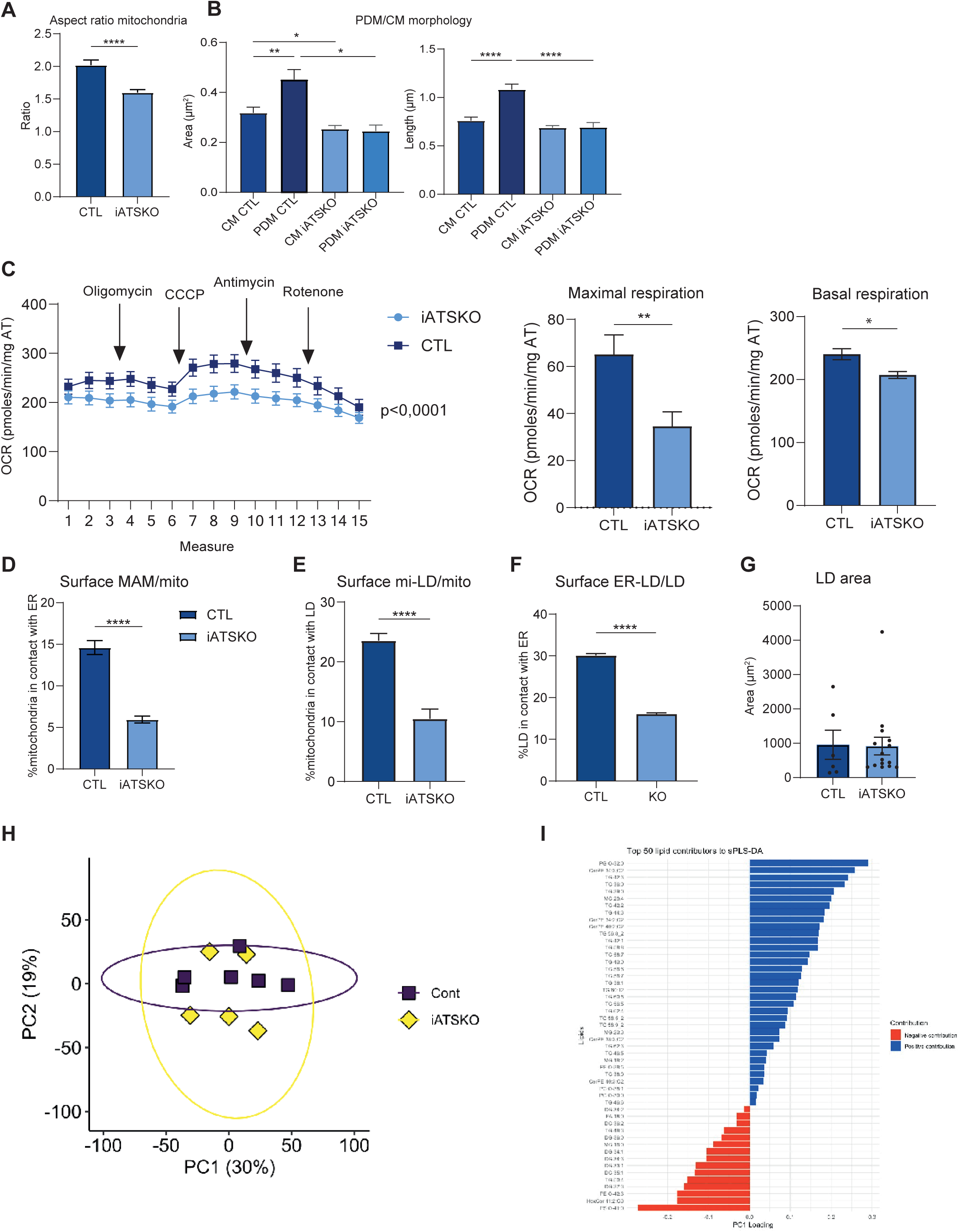
Inguinal adipose tissue was collected from Bscl2^lox/lox^ (Control) and Bscl2^lox/lox^; ERT2-Adipoq-Cre (iATSKO) mice in the random-fed state, 14 days after tamoxifen injection (100 mg/kg). TEM was used to quantify mitochondria aspect ratio (A), area and length (B). Seahorse analysis was performed on 2 mg explants, with three biopsies per mouse (C). TEM was used to quantify the following organelle contacts: ER–mitochondria (MAM), normalized to the surface of mitochondria (D); mitochondria–lipid droplet (Mi–LD) contacts, normalized to the surface of mitochondria (E); and ER–lipid droplet (ER–LD) contacts, normalized to lipid droplet (LD) surface area (F). LD surface area was also quantified (G). LDs were isolated from iATSKO and control mice, and proteomic (H) and lipidomic (I) analyses were performed. sPLDA analysis identified the lipids that contributed the most to the separation of the genotype profiles (I).

**Supplemental 2.**
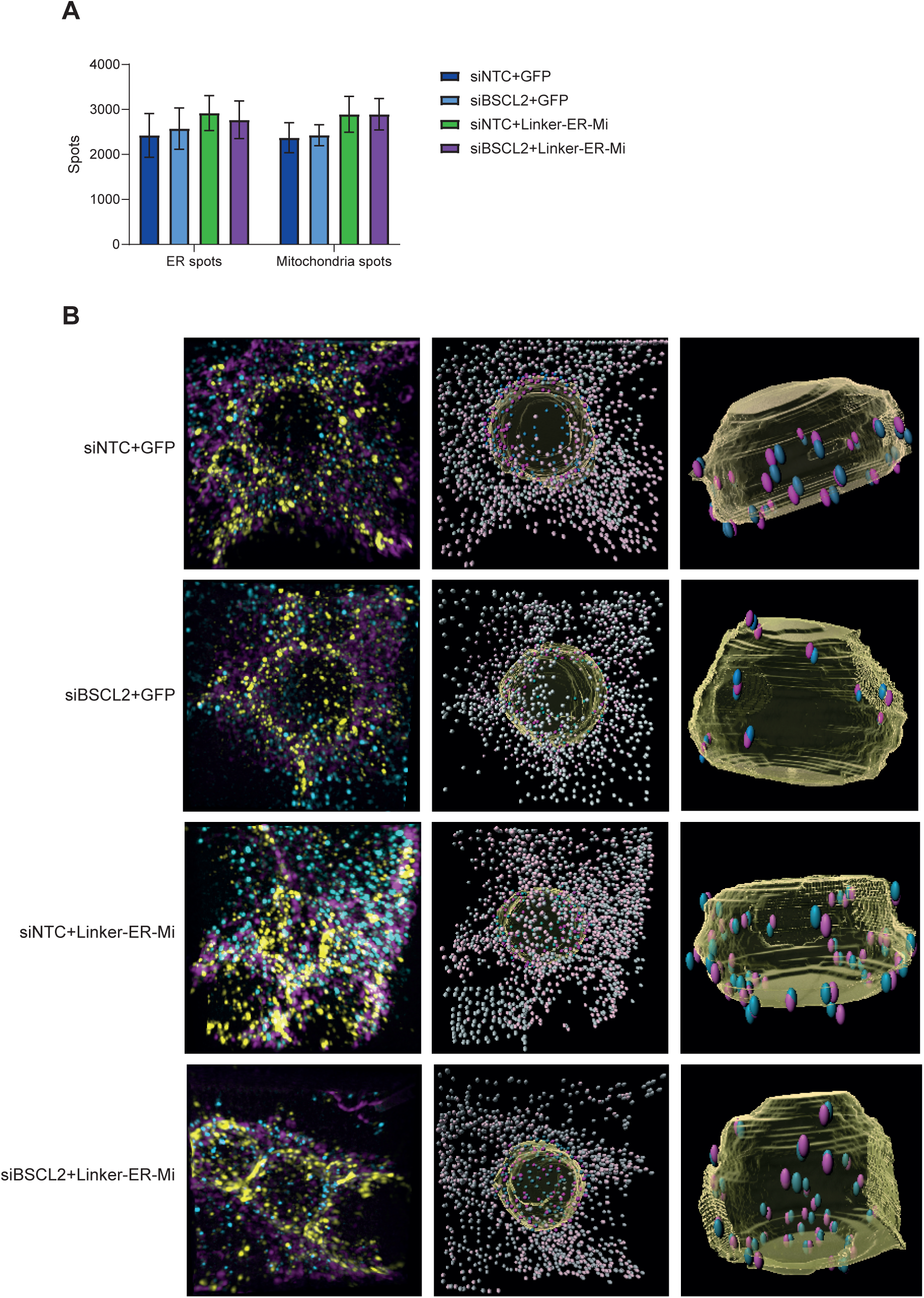
Structured Illumination Microscopy (SIM) was used to visualize mitochondria (Tom20), endoplasmic reticulum (PDI), and lipid droplets (PLIN1). Spatial proximity analyses were conducted using Imaris software, and the total number of spots was quantified (A). Representative images are shown (B).

**Supplemental 3.**
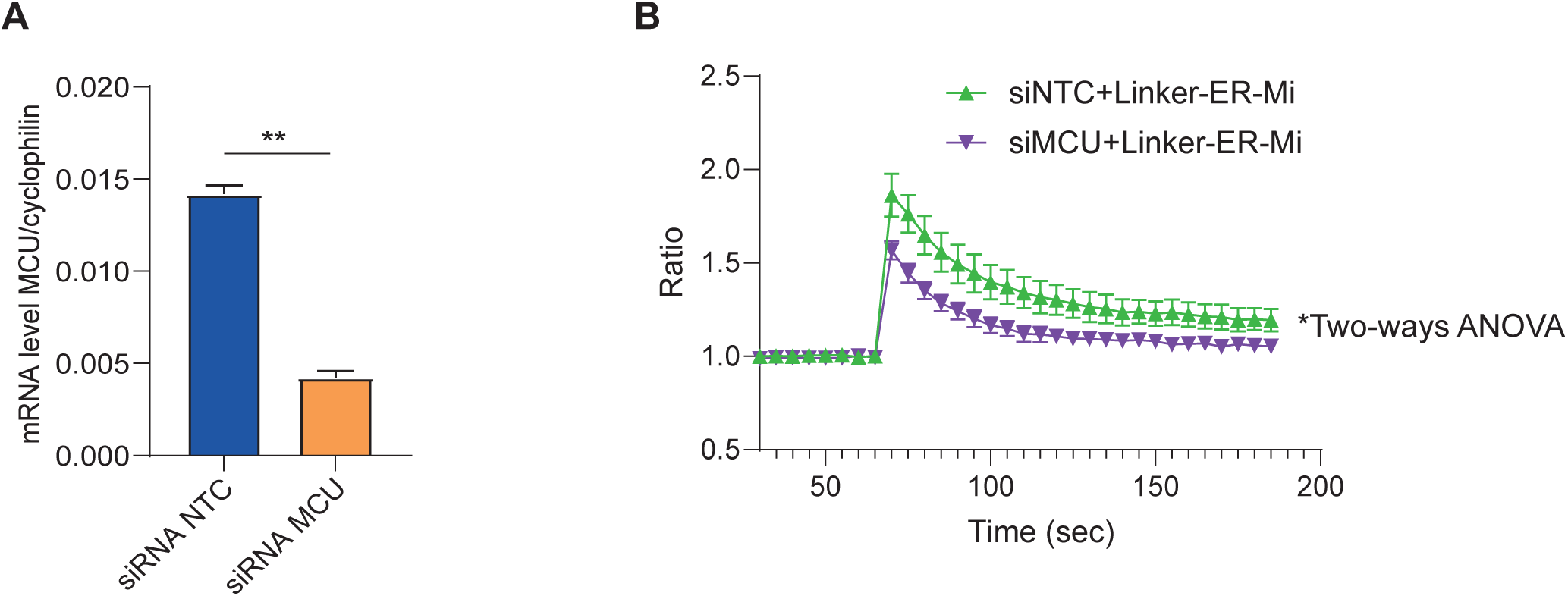
3T3-L1 adipocytes were reverse-transfected with the indicated siRNAs and infected with adenovirus expressing either GFP or Linker-ER-Mi. MCU mRNA levels were quantified by qPCR (A). Mitochondrial calcium was monitored using caged IP₃ and Rhod2. After 60 seconds of baseline acquisition, UV flash photolysis was used to release IP₃. The Y-axis represents the fluorescence ratio relative to baseline (B).

**Supplemental 4.**
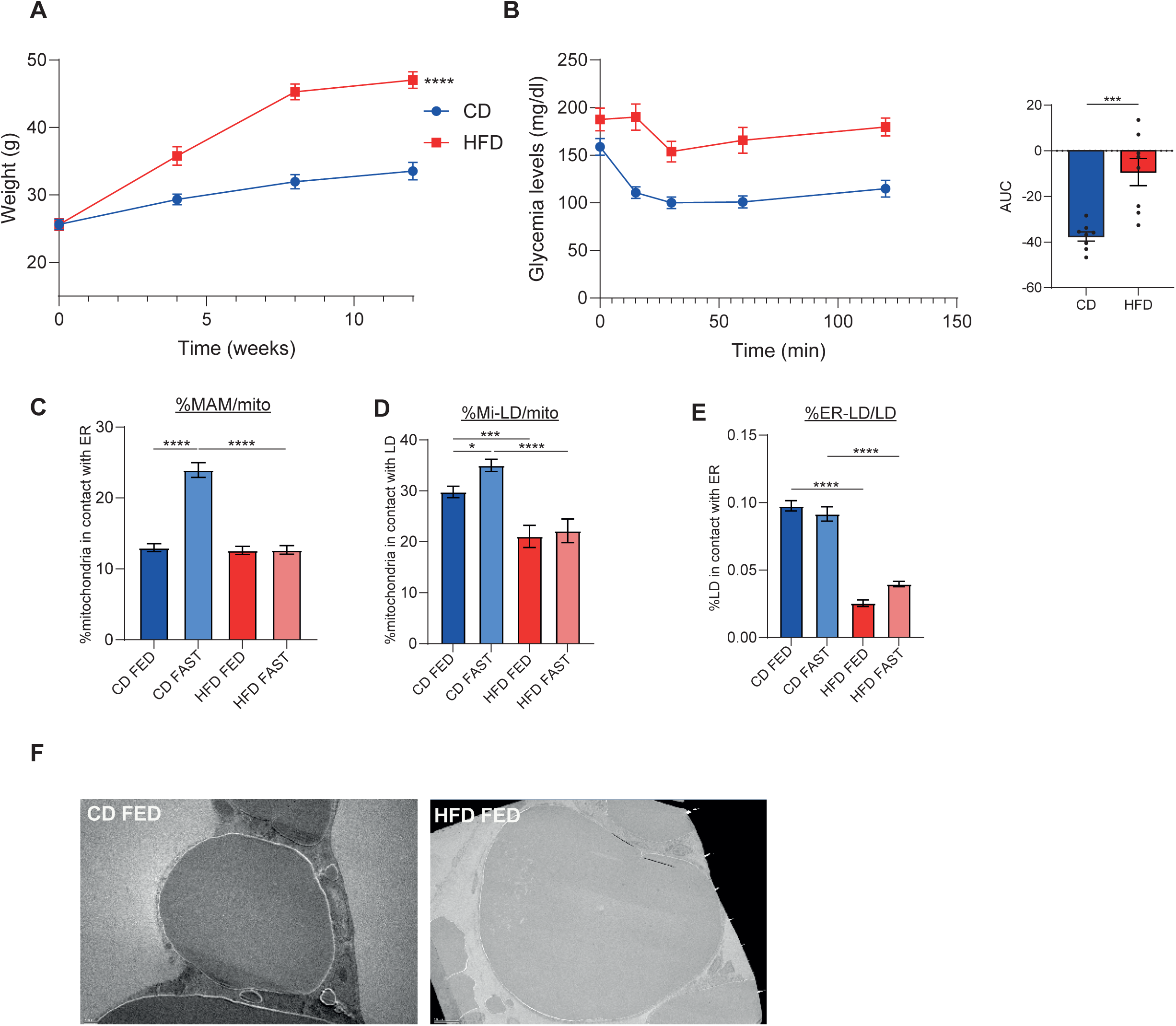
Mice were fed chow or high fat diet for 3 months. Weight was monitored (A) and insulin sensitivity was evaluated after 12 weeks of diet (B). Gonadal adipose tissue from mice fed chow diet or high fat diet were collected in fed or 18-hour fasted state. Transmission electron microscopy was used to quantify the following organelle contacts length with FIJI: ER-mitochondria contact (MAM, C), mitochondria-LD contact (Mi-LD, D), and ER-LD (E). Histological images of gonadal adipose tissue (F). Bars: mean +/- SEM, n=3 mice per group, 5 cells per mouse. *p<0.05, ***p<0.001, ****p<0.0001, Mann-Whitney test.

**Supplemental 5.**
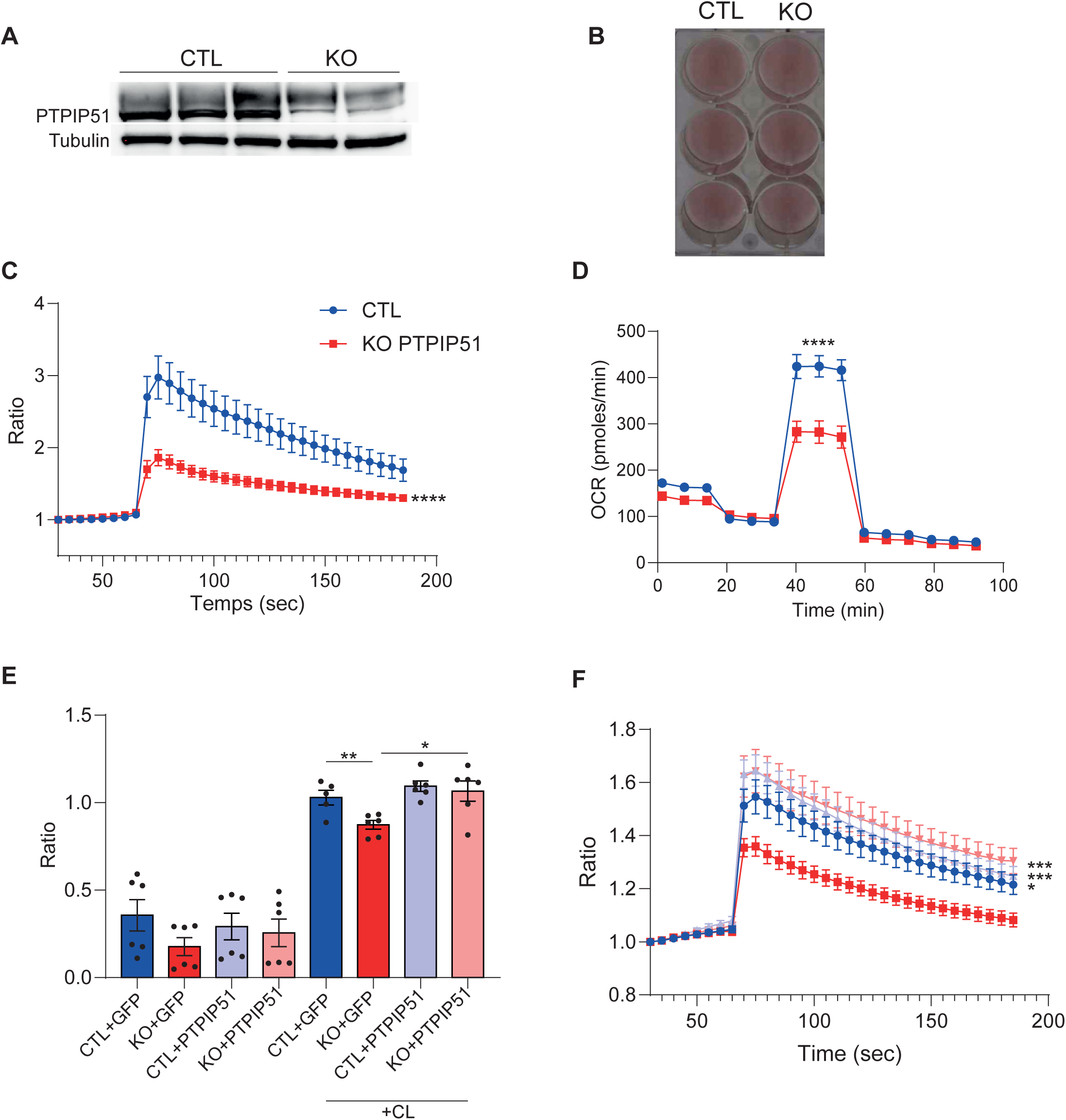
Western Blot image in control and KO PTPIP51 3T3-L1 adipocytes (A). Oil Red O staining of CTL and KO PTPIP51 3T3-L1 cells (B). Mitochondrial calcium was monitored using caged IP₃ and Rhod2. After 60 seconds of baseline acquisition, UV flash photolysis was used to release IP₃. The Y-axis represents the fluorescence ratio relative to baseline (C). OCR was measured using the seahorse system (D). Lipolysis was measured by quantifying glycerol release in extracellular media after 2 hours of incubation +/-CL316-243 (E). Mitochondrial calcium was monitored as mentioned above (F). Bars: mean +/- SEM, each condition in triplicate, n=3 experiments, *p<0.05, ***p<0.001, ****p<0.0001, Mann-Whitney test.

**Supplementary Videos 1 and 2** show 3D reconstructions from FIB-SEM acquisitions of adipose tissue sections from control (Video 1) and iATSKO (Video 2) mice. Each animation combines aligned raw image stacks with organelle segmentation, enabling visualization of the spatial organization of lipid droplets and mitochondria within the tissue

**Supplementary Videos 3 and 4** show 3D reconstructions from FIB-SEM acquisitions of 3T3-L1 adipocytes treated with NTC control (Video 3) or siBSCL2 (Video 3) si RNA. Each animation combines aligned raw image stacks with organelle segmentation, enabling visualization of the spatial organization of lipid droplets and mitochondria within the adipocytes

## Materials and Methods

### 3T3-L1 Cell Culture and Differentiation

3T3-L1 cells were maintained in medium A (DMEM with 4.5 g/L glucose, 10% calf serum, 1% penicillin/streptomycin, and 1% glutamine). Cells were seeded at a density of 20,000 cells/mL in medium B (DMEM with 4.5 g/L glucose, 10% fetal calf serum (FCS), 1% penicillin/streptomycin, and 1% glutamine). Two days post-confluence, adipogenic differentiation was induced using medium B supplemented with 2 µM insulin, 500 µM IBMX, and 1 µM dexamethasone (differentiation medium, DMI). On day 3, cells were switched to medium B containing only 2 µM insulin. From day 6 onward, cells were maintained in medium B alone.

### Animals

Male C57BL/6 mice were fed a high-fat diet (HFD) (61% of calories from fat, 20% of calories from protein, and 20% calories from carbohydrate, #D12492 Safe Diets), or a normal chow diet *ad libitum* for a period of 12 weeks. All mice had *ad libitum* access to food and water and were housed in the same open mouse facility on a 12/12-h light/dark day/night cycle. Animal care and study protocols were approved by the French Ministry of Education and Research and the ethics committee N°6 in animal experimentation and were in accordance with the EU Directive 2010/63/EU for animal experiments.

### Insulin tolerance test (ITT)

Food was removed 2 hours before the initiation of the tolerance test. For ITT, mice were injected i.p. with 1 IU/kg body weight insulin at time0. Blood glucose levels were monitored using glucometer strips at 0, 15, 30, 60 and 120 min on 2.5µL samples collected from the tail.

### Generation of PTPIP51 Knockout Cells in 3T3-L1 Line

3T3-L1 preadipocytes were cultured in stromal vascular fraction (SVF) medium and passaged upon reaching ∼50% confluence. Cells were trypsinized, collected, and counted. A total of 5.10^5^ cells were electroporated using the Amaxa Cell Line Nucleofector Kit V (VCA-1003, Lonza) with a ribonucleoprotein complex consisting of recombinant Cas9 protein (Alt-R™ S.p. Cas9 Nuclease V3, 21µM final, IDT), and two pre-assembled crRNA:tracrRNA duplexes (crRNA sequences ACCAAGUAACAGUCCCAACC and UAUGACCUGUGCUCUUACCC, IDT), targeting the *murine Rmdn3* gene (encoding PTPIP51, exon1). As a control, cells were electroporated with Cas9 and tracrRNA only, without crRNA. Forty-eight hours post-electroporation, cells were trypsinized and sorted using flow cytometry (FACS Aria III/Influx), with single cells seeded into individual wells of a 96-well plate to generate clonal populations. Clones were screened for successful biallelic gene disruption using genomic PCR (PTPIP51-F: *GGTTGTCACGTACAGGGTGT ;* PTPIP51-R : *GGCGCGACCAGAACTAAAAG*). Selected clones were expanded and evaluated for their adipogenic differentiation potential. Respectively four and five clones showing comparable differentiation capacities were pooled to generate two distinct populations referred to as “Control” and “PTPIP51 KO” throughout this study.

### siRNA Transfection and Reverse Adenoviral Infection

On day 6 of differentiation, cells were harvested from T75 flasks using TrypLE Express (ThermoFisher), and resuspended in 12 mL of medium B per flask. For transfection, a mix containing 20 nM siRNA, 2.5 µL Lipofectamine RNAiMAX (ThermoFisher), and 47.5 µL Opti-MEM (ThermoFisher) was prepared and added to the bottom of collagen IV-coated Ibidi 8-well chambers. Then, 0.25 mL of the cell suspension was added to each well. When required, adenoviruses were added directly to the transfection mix at the indicated MOI (see figure legends). Cells were harvested and analyzed 72 h post-transfection. The siRNA pool targeting BSCL2 was obtained from Horizon Discovery.

### Proximity Ligation Assay (PLA)

PLA was performed on 3T3-L1 cells using the Duolink™ In Situ FarRed kit (#DUO92013, Sigma-Aldrich) following the manufacturer’s instructions. After PBS washes, cells were fixed in 4% PFA for 10 minutes and washed again in PBS. Permeabilization was performed using 0.2% saponin in PBS. Blocking was done at room temperature for 60 minutes using a solution of 3% BSA, 0.1% saponin, and 20 mM glycine in PBS. Primary antibodies were incubated overnight at 4°C in PBS containing 2% BSA, 0.1% saponin, and 20 mM glycine. Antibodies used: mouse anti-IP3R1-2-3 (1:100, Santa Cruz 377518), mouse anti-Perilipin1 (1:500, Progen 651156), rabbit anti-VDAC1 (1:200, Abcam 15895), rabbit anti-VAPB (1:200, Sigma-Aldrich HPA013144). Subsequent steps followed the kit protocol. Nuclei were counterstained with DAPI and washed in Tris-HCl (200 mM), NaCl (137 mM), and 0.05% Tween. Z-stack images were acquired on a Nikon A1RSi confocal microscope with a 60x oil immersion objective. PLA focus quantification per cell was performed using ImageJ 2.0 with the following settings: nuclear area 25–100 µm²; PLA dot size 0–1 µm². Statistical analysis was conducted using the Mann–Whitney test (Prism 7.0).

### Fluorescence Recovery After Photobleaching (FRAP)

Differentiated 3T3-L1 adipocytes were incubated for 4 h in medium B supplemented with BODIPY558/568 C12 (2 µM) and oleic acid (OA, 20 µM). After PBS washes, cells were transferred to FluoroBrite medium containing OA (20 µM). Imaging was performed on a Nikon A1 confocal microscope using a 20x objective. A region of interest (ROI) located on a lipid droplet (LD) was selected, and fluorescence was recorded for 10 seconds before photobleaching with a high-intensity laser for 500 ms. Fluorescence recovery was monitored for 90 seconds. Data were analyzed using ImageJ 2.0 and EasyFRAP.^48^ Two-way ANOVA was used for statistical comparisons (Prism 10).

### Super-resolution Structured Illumination Microscopy (SIM)

Differentiated adipocytes treated with oleate (700µM) during 6 hours were washed twice with PBS, fixed in 4% PFA for 10 minutes, and permeabilized using 0.5% saponin for 20 minutes. Blocking was done at room temperature for 60 min using a solution of PBS-Tween BSA 3%. Primary antibodies were incubated for 1 h at room temperature in PBS with 1% BSA, followed by three PBS washes. Antibodies used: mouse anti-PDI (1:50 Thermo MA3-018), guinea pig anti-Perilipin1 (1:50 Progen), rabbit anti-TOM20 (1:100 Thermo MA5-34964). Secondary antibodies were incubated 30 min under the same conditions. Slides were mounted using ProLong mounting medium. SIM imaging was performed using a Nikon A1 N-SIM microscope equipped with a 100×/1.40 NA oil immersion objective. Structured illumination patterns were projected using 488, 561, and 642 nm lasers. Fluorescence was captured using an EMCCD camera (iXon 885, Andor Technology). 3D stacks were reconstructed with NIS software. LD segmentation was performed using a custom workflow based on a 2D retrained model Cellpose model for 2D segmentation^49^, and 3D reconstruction using CellStitch ^50^ (available upon request) and imported afterwards in IMARIS, where spot segmentation and distance assessment (spot to surface and spot to spot) were then carried out.

### FIB-SEM sample preparation

Fat tissues from wild-type (WT) and mutant specimens were dissected and washed three times in phosphate-buffered saline (PBS). Samples were then fixed in 2,5% glutaraldehyde (Electron Microscopy Sciences, MA, USA) prepared in 0,2M cacodylate buffer (pH 7.2) overnight at 4°c. Following fixation, tissues were washed three times in fresh 0,1M cacodylate buffer (pH 7.2) and post-fixed in 1% osmium tetroxide (Electron Microscopy Sciences) in 0,1M cacodylate buffer supplemented with 1,5% potassium ferrocyanide for 1h in the dark. After three washes in distilled water, samples were incubated for 30 min at room temperature in 0,2% tannic acid (in water), then post-fixed a second time in 1% osmium tetroxide for 1h and washed again. Samples were dehydrated through a graded ethanol series (25%, 50%, 70%, 90% and 100%) and embedded in epoxy resin (PolyBed 812, Electron Microscopy Sciences), followed by polymerization for 58h at 60°c.

### FIB-SEM imaging

For Focused Ion Beam-Scanning Electron Microscopy (FIB-SEM) acquisition, resin-embedded tissue blocks were mounted on aluminum stubs and sputter-coated with 20 nm gold-palladium layer using a Leica ACE600 coater. Imaging was performed with the specimen stage titled at 54°c, maintaining a 5 nm working distance at the coincidence point of the electron and gallium beams. Serial block milling was performed using a 500 pA milling current, removing 16 nm slices from the specimen surface at each step. Images were acquired at 1.5 kV using the in-lens EsB detector, with the EsB grid set between -300V and -500V. The final dataset had a voxel size of 15nm in x, y and z dimensions.

### FIB-SEM image analysis

Images were aligned with the plug-in Stackreg from Fiji : P.Thévenaz, U.E.Ruttimann, M.Unser, “A Pyramid Approach to Subpixel Registration Based on Intensity” IEEE Transactions on Image Processing, vol.7, no.1, pp.27-41, January 1998. The rest of Workflow was done on Amira software version 2024.2. The segmentations were produced with the AI Assisted Segmentation module. 7 ROIs (rectangular regions containing the annotations and the background given the neural network) were delimited where the labels mitochondria, the adipocyte plasmic membrane, the adipocyte nucleus and the lipid droplets were manually segmented for the wild-type condition. Whereas 10 ROIs were produced for the labels mitochondria, the adipocyte plasmic membrane and the lipid droplets segmentation for the CRE condition. In both cases the annotations were manually done with the brush or the magic wand tools. Then the ResNet18 deep learning model was selected with 30 epochs chosen for the training. In the end the prediction was corrected manually either in 2D with the brush, the pick or the lasso tools, or in 3D with the pick or the lasso tools.

### Mitochondrial Calcium Flux Assay

Differentiated adipocytes were washed with HBSS and incubated for 1 h at 37°C in a loading solution (900 µL HBSS + 100 µL Fluoforte reagent B, ENZO kit) containing 3 µM caged IP3 (Sichem), 5 µM Fluo4 (Thermo Fisher) or 10 µM Rhod2 (Thermo Fisher), and 0.3% Pluronic F-127 (Thermo Fisher). After loading, cells were washed in HBSS. Time-lapse imaging was performed using a Leica DMI 6000B microscope controlled by Metamorph software. Following baseline acquisition, IP3 photorelease was triggered using a UV flash.

### Lipidomics Analysis

Differentiated adipocytes were resuspended in 400 µL deionized water. Lipid extraction was performed by adding 1600 µL of cyclohexane/isopropanol (3:2), followed by 1 h agitation at room temperature and centrifugation at 10,000 g for 5 min. The organic upper phase was transferred to glass tubes and evaporated at 60°C under nitrogen. Dried lipids were resuspended in isopropanol/acetonitrile/water (65:30:5) and analyzed by liquid chromatography coupled with mass spectrometry (LC-MS).

### Fluxomics Analysis

Metabolites were analyzed by LC-HRMS. Cells were incubated for 24 h in serum-free DMEM lacking glutamine, pyruvate, and phenol red, supplemented with 25 mM 13C6-glucose (Eurisotop CLM-1396-1). After collection, cells were lysed in 200 µL absolute methanol and centrifuged at 12,000 rpm for 15 min. Supernatants were dried under nitrogen at 45°C and resuspended in water containing 0.1% formic acid. Samples were analyzed using a Synapt G2 HRMS Q-TOF mass spectrometer (Waters) in negative ion mode, with separation on an Acquity H-Class UPLC system. Ionization parameters included a 1 kV capillary voltage, 30 V cone voltage, 900 L/h desolvation gas flow, and 550°C temperature. Data were processed with MassLynx, TargetLynx, and IsoCor. Metabolite levels were normalized to cell number (per 10⁶ cells). 13C enrichment was calculated as the ratio of labeled to unlabeled metabolites. Mann–Whitney tests were used for statistical analysis (GraphPad Prism).

### Transmission Electron Microscopy (TEM)

Adipose tissue (1 mm³) or differentiated adipocytes were fixed for 16 h at 4°C in 2.5% glutaraldehyde (LFG Distribution) in 0.1 M Sorensen buffer (pH 7.4). After rinsing in 0.2 M cacodylate buffer (pH 7.4), samples were post-fixed in 2% osmium tetroxide/1.5% potassium ferrocyanide for 45 min at room temperature. Samples were dehydrated in a graded ethanol series and embedded in EMbed 812 resin (LFG Distribution), polymerized at 60°C for 48 h. Semi-thin (1 µm) sections were stained with methylene blue/Azur II and observed under an Olympus AX-60 light microscope. Ultrathin sections (60 nm) were examined with a Jeol JEM 1400 TEM operating at 120 keV.

### Western Blotting

Cells or tissues were lysed in buffer containing 1% Igepal CA-630, 0.05–0.5% sodium deoxycholate, 0.1% SDS, 250 mM Tris-HCl (pH 7.5), and 150 mM NaCl, supplemented with protease inhibitors. Lysates were centrifuged at 13,000 g for 10 min at 4°C. Equal amounts of protein were resolved by SDS-PAGE, transferred to membranes, and blocked in 5% milk in TBS-T (0.1% Tween-20) for 1 h at room temperature. Membranes were incubated with primary antibodies overnight at 4°C and with HRP-conjugated secondary antibodies for 1 h at room temperature. Detection was performed using ECL substrate and visualized with a ChemiDoc system (Bio-Rad). Band intensities were quantified using ImageJ FIJI and normalized to total protein content (Stain-Free system, Bio-Rad).

### Statistical Analysis

Data are expressed as mean ± SEM. Parametric or non-parametric tests are specified in figure legends. Area under the curve (AUC) values were compared using unpaired t-tests. Two-way ANOVA was used where indicated to test for interactions between experimental groups and variables. All analyses were performed using GraphPad Prism.

